# Parsimonious EBM: generalising the event-based model of disease progression for simultaneous events

**DOI:** 10.1101/2022.07.10.499471

**Authors:** CS Parker, NP Oxtoby, AL Young, Alzheimer’s Disease Neuroimaging Initiative, DC Alexander, H Zhang

## Abstract

This study introduces the parsimonious event-based model of disease progression (P-EBM). The P-EBM generalises the event-based model of disease progression (EBM) to allow inference of fewer disease progression stages than the number of input biomarkers. The original EBM is designed to estimate a single distinct biomarker abnormality, termed an event, at each model stage. By allowing multiple events within a common stage, the P-EBM prevents redundant complexity to permit discovery of parsimonious sequences of disease progression - those that contain purely serial events, as in the original EBM, as well as those containing one or more sets of simultaneous events. This study describes P-EBM theory, evaluates its sequence estimation and staging performance and demonstrates its application to derive a parsimonious sequence of disease progression in sporadic Alzheimer’s disease (AD). Results show that the P-EBM can accurately recover a wider range of sequences than EBM under a range of realistic experimental scenarios, including different numbers of simultaneous events, biomarker disease signals and dataset sizes. The P-EBM sequence successfully highlights redundant biomarkers and stages subjects using fewer biomarkers. In sporadic AD, the P-EBM estimates a shorter sequence than the EBM with substantially higher likelihood which plausibly suggests that some biomarker events appear simultaneously. The P-EBM has potential application for generating new insights into disease evolution and for suggesting efficient biomarker collection strategies for patient staging.

## 1. Introduction

Understanding the temporal evolution of disease is crucial for the development of effective treatments. Imaging and non-imaging biomarkers provide a window into the underlying pathophysiology and clinical manifestation of disease through the measurement of observable disease phenomenon. Estimating the temporal trajectory of biomarker abnormalities therefore informs on disease temporal evolution, with potential to aid personalised prognosis through patient staging and to facilitate development of interventions by providing insights into disease mechanisms.

Sequences of biomarker progression have been previously derived using neuropathological or hypothetical approaches. For example, the Braak and Braak staging system describes the spatiotemporal evolution of tau pathology in AD based on pathological examination of post-mortem brain tissue (Braak, H & Braak, E 1991). The “Jack curves” describe the hypothetical dynamic evolution of AD biomarkers based on review of research studies and clinical findings (Jack Jr. et. al. 2010, 2013). While informative, these approaches rely on qualitative labour-intensive evaluation, which makes them to some degree subjective, and difficult to apply to new biomarker datasets.

To overcome this, disease progression modelling approaches have been developed which utilise large datasets for objective and data-driven inference of disease progression sequences (Green et. al. 2011, Young et. al. 2018, Oxtoby et. al. 2017, Young et. al. 2024). In the ideal scenario, the temporal trajectory of different biomarkers is inferred from longitudinal subject data acquired throughout the disease course. Yet in practice, typically only cross-sectional or short-term longitudinal biomarker data are available. There is therefore a need to estimate disease progression from such data. The event-based model of disease progression (EBM) is a data-driven approach that estimates the evolution of biomarker abnormalities from cross-sectional or short-term longitudinal data (Fonteijn et. al. 2012). The EBM has been applied to a variety of diseases, including AD (Fonteijn et. al. 2012, Huang & Alexander 2012, Young et. al. 2014, Oxtoby et. al. 2018, Venkatraghavan et. al. 2017, Young et. al. 2018, Firth et. al. 2020, Young et. al. 2021, Vogel et. al. 2021, Wijeratne et. al. 2023, Young et. al. 2023), Huntington’s disease (Fonteijn et. al. 2012, Wijeratne et. al. 2018), Parkinson’s disease (Oxtoby et. al., 2021), multiple sclerosis (Eshaghi et. al. 2018, 2021) and amyotrophic lateral sclerosis (Gabel et. al. 2020).

The EBM represents the temporal progression of disease biomarkers as a series of stages whereby each biomarker undergoes a transition, termed an event, from a normal state to an abnormal state. A key modelling choice of the original EBM (Fonteijn et. al. 2012), as well as its subsequent variants (Huang & Alexander 2012, Venkatraghavan et. al. 2018, Young et. al. 2018, Firth et. al. 2020, Young et. al. 2021, Tandon et. al. 2023, Young et. al. 2023), is that biomarker events are ordered serially, i.e., no two events may occur at the same modelled disease stage. However, this assumption may not always hold - events may be better modelled as simultaneous when assigned to a single disease stage. This is the case when their temporal trajectories are simultaneous, or when there is a temporal difference in their trajectories that is unresolvable in practice (i.e., when temporal sampling is coarse relative to their trajectory differences). Such apparently simultaneous events cannot be inferred by the EBM as they are excluded from the model by construction. The positional uncertainty that the EBM estimates may suggest the presence of simultaneous events, but can also simply reflect a high biomarker variance.

The serial events assumption adopted by EBM forces an equality between the number of modelled disease stages and the number of biomarker events. By not accounting for simultaneous events, the EBM may artificially enforce a serial event ordering, thereby over-estimating the number of disease stages. This has potentially adverse ramifications for disease understanding and for patient staging.

To overcome this, we introduce the parsimonious event-based model of disease progression (P-EBM), a generalisation of the EBM that can estimate sequences containing simultaneous events. By allowing simultaneous events, a wide range of parsimonious disease progression sequences can be estimated from a given biomarker data input. Such sequences inform on disease evolution and highlight the key biomarker measures needed for patient staging. This study demonstrates the P-EBM by generalising the original EBM (Fonteijn et. al. 2014) to allow inference of simultaneous (multi-biomarker) events within a single disease stage. We introduce the theory behind P-EBM and evaluate its performance against the EBM in simulated and real data. P-EBM is applied to reduce biomarker collection requirements for patient staging and to infer a parsimonious sequence of imaging and non-imaging biomarker events in sporadic AD.

## 2. Theory

### 2.1. Overview of the event-based model of disease progression

The EBM predicts biomarker measurements of a study cohort from a sequence of biomarker abnormalities (Fonteijn et. al. 2012). The sequence encodes a temporal order of events - the order in which each biomarker transitions from a normal state to an abnormal state. The inversion of this model, typically performed through maximum likelihood estimation, allows inferring the sequence from a set of biomarker measurements. An overview of the EBM, highlighting the role of the likelihood and sequence estimation, is shown in Fig. 1.A.

**Figure 1.**
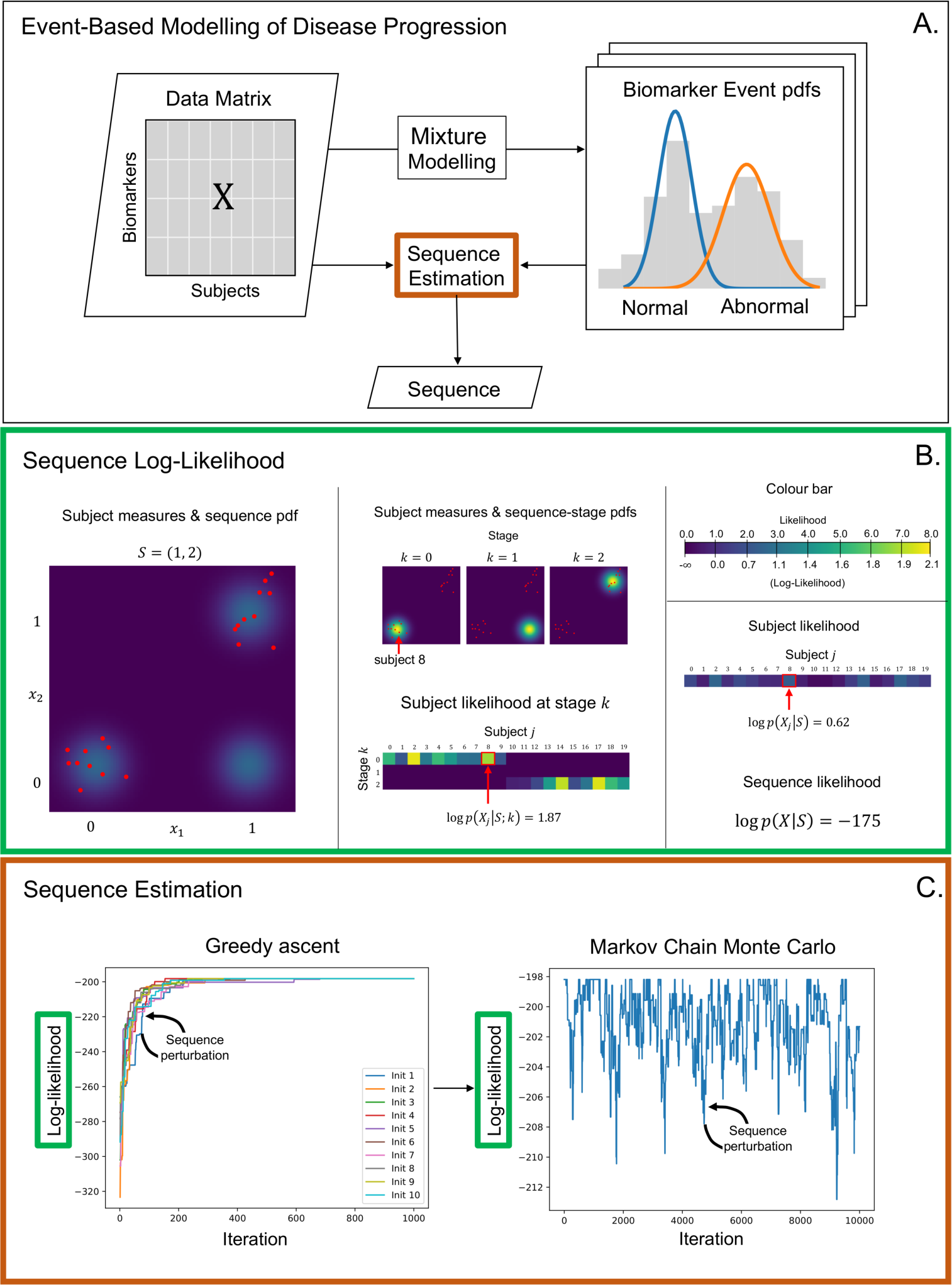
Overview of the EBM highlighting the roles of likelihood calculation and sequence estimation. The EBM infers a sequence of biomarker events from a data matrix of cross-sectional biomarker measurements (**A**). To do this, EBM uses the pre- and post-event biomarker distributions (“Normal” and “Abnormal”) to evaluate the likelihood of a proposed sequence (**B**) and estimates the maximum likelihood sequence using greedy ascent and/or MCMC (**C**). Panel (**B**) shows an example sequence likelihood calculation for the case of two biomarkers with the proposed sequence S = (1,2) and given example subject measures comprising a dataset X. In (**B**) subject measures are plotted over probability density functions (pdf’s), which visualise p*X_(·)_+S, (the sequence pdf) and p*X_(·)_+S; k, (the sequence stage pdf) across the space of unobserved subject data X_(·)_. For a given dataset of subject measures (red dots), the subject likelihood at stage k, subject likelihood, and sequence likelihood, are shown in middle and right of (**B**).

The foremost assumption of EBM is that each biomarker undergoes an irreversible transition from a normal (pre-event) state to an abnormal (post-event) state. Other key assumptions are (i) subjects are sampled from the same disease trajectory; (ii) biomarker measurements for a subject at a particular stage in the sequence are independent of one another; (iii) the set of biomarker measurements for each subject is independent from other subjects.

Given the input data X, an N-by-J matrix containing N biomarker measurements for J subjects (Fig. 1.A), and a proposal sequence S, the sequence likelihood p(X|S), under assumption (iii), is the product of subject-wise sequence likelihoods p(X_!_|S):

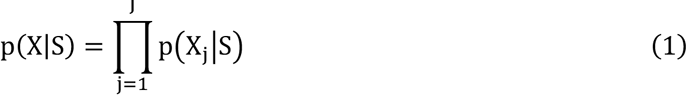

where X_!_ is a column of X corresponding to the N biomarker measurements for subject j (for instance a red dot in Fig. 1.B). Next, we describe the sequence representation and subject-wise likelihood (subject likelihood henceforth) formulation for the EBM, which assumes events occur in series. Afterwards, we generalise the sequence and likelihood for simultaneous events.

### 2.2. Sequence and subject likelihood

#### 2.2.1. Event-based model of disease progression

The assumption that events are strictly serial means that the sequence, S, is a permutation of the biomarker indices {1, … , N}. Each element, S(i), holds the biomarker event occurring at the i‘th disease stage. For example, for a sequence of four biomarkers S = (2,3,4,1), the first event (disease model stage 1) occurs for biomarker 2, followed by biomarker 3, then biomarker 4 and finally biomarker 1. At a disease model stage of 0, no events have yet occurred.

Assuming independence of biomarker measurements for a subject at a particular stage k (assumption (ii)), the subject likelihood at stage k, p(X_!_|S, k), is the product of probabilities for observing each biomarker measurement given the events have occurred for biomarkers {S(i) | i = 1, … k} but have not occurred for biomarkers {S(i) | i = k + 1 , … , N}:

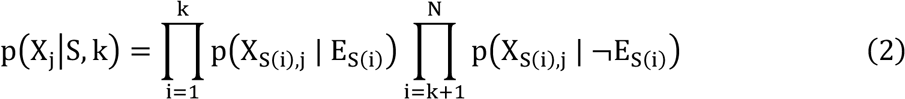

where E_%(’)_ denotes the event has occurred for biomarker S(i) and ¬E_%(’)_ that it has not. As each subject’s stage is unknown a priori, the subject likelihood is calculated as a marginalisation over all disease stages:

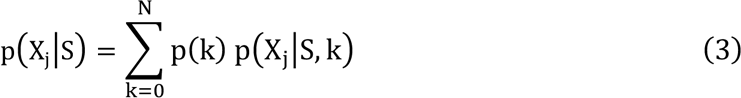

A non-informative (uniform) prior probability of each stage is typically assumed, 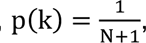 where N + 1 (or equivalently |S| + 1) is the number of stages in the sequence. By substituting Eq. (2) into Eq. (3), the sequence likelihood as defined in Eq. (1) can be written as:

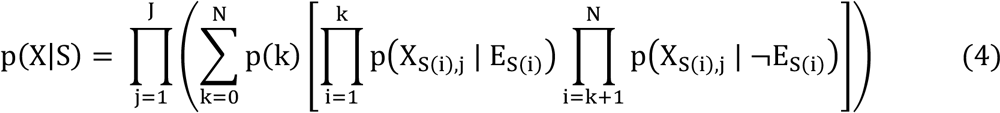

An example sequence likelihood calculation for the case of two biomarkers is shown in Fig. 1.B. As the EBM sequence contains only one biomarker event at each model stage, it can only evaluate the likelihood of sequences of serial events.

#### 2.2.2. Parsimonious event-based model of disease progression

To generalise the EBM for simultaneous events, the sequence specification is updated from an ordered list of biomarker indices to an ordered list of sets. Each set, S(i), contains one or more biomarker indices, corresponding to the events at disease stage i. For example, a serial sequence of four biomarkers could be S = ({2}, {1}, {3}, {4}) and a sequence containing simultaneous events could be S = ({2}, {1, 3}, {4}).

The subject likelihood at stage k, p(X_!_|S, k), can be computed as before, but this time referencing the P-EBM sequence, where the events have occurred for biomarkers ⋃_$,’,-_ S(i), but have not occurred for biomarkers ⋃_-.’,|%|_ S(i). Hence, the subject’s likelihood given their stage is:

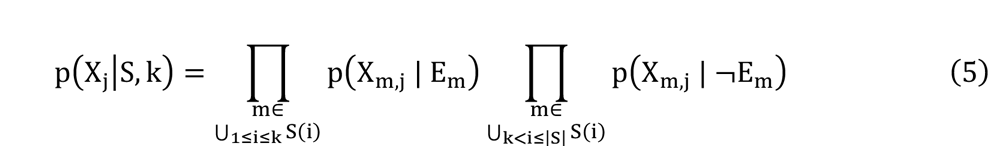

The unknown subject’s stage is again marginalised to give the subject’s likelihood. But given the length of the S-EBM sequence can vary, the number of stages is |S| + 1, making the non-informative prior probability of each stage 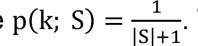. The subject’s likelihood given their stage is unknown a priori is then written as:

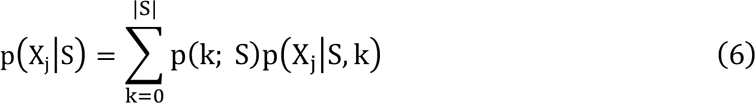

By substituting Eq. (5) into (6) and then Eq. (6) into (1), the P-EBM sequence likelihood is:

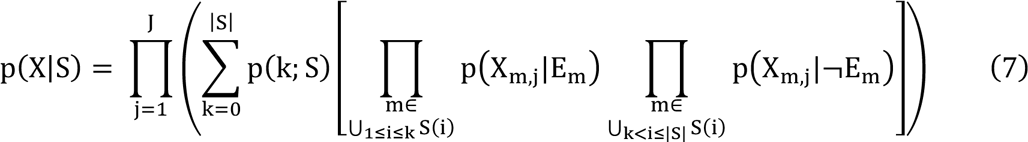

For sequences consisting of only serial events, the likelihood is equal to the EBM likelihood in Eq. (4). This more general sequence representation and associated likelihood can used to evaluate a wider range of proposed sequences than the EBM, including shorter sequences with fewer stages where one or more single-biomarker events occur simultaneously, as a multi-biomarker event. Fig. 2 shows an example of the P-EBM’s capability to estimate sequences containing either serial or simultaneous events using the likelihood formulation in Eq. (7).

**Figure 2.**
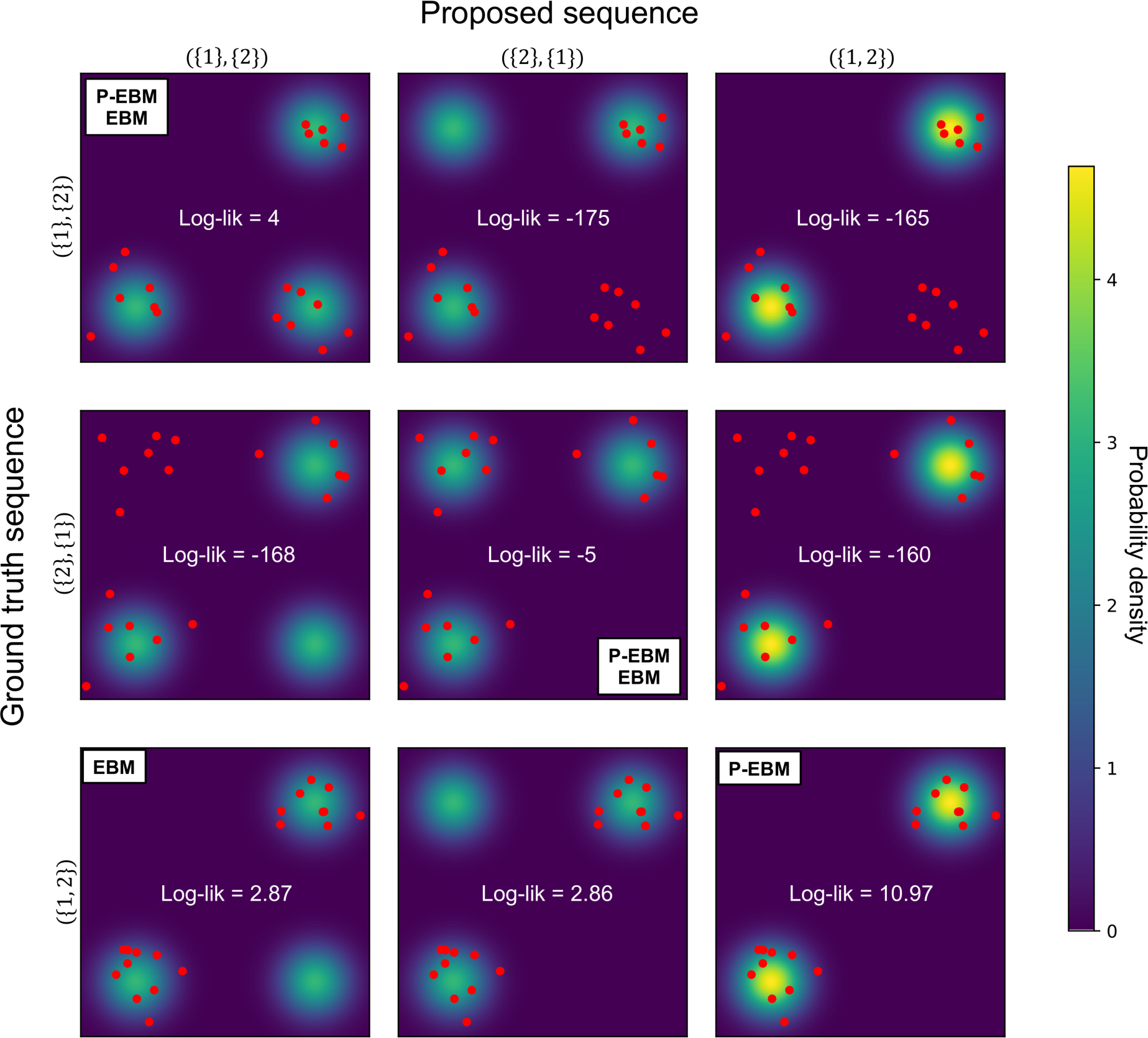
Likelihoods of proposed sequences for different datasets. Colour maps show the pdf p(X_(·)_|S) for the three possible proposal sequences of two biomarkers (columns). Overlaid in red dots is a dataset of twenty subjects, generated under one of the same three sequences, with each row plotting the same dataset for a given ground truth sequence. For each dataset, the proposed sequence estimated by the P-EBM and EBM is denoted by the text inset. C-EBM and P-EBM both correctly infer serial sequences (rows 1 and 2), but only P-EBM can infer sequences of simultaneous events (row 3). C-EBM incorrectly estimates a serial sequence for data generated from a simultaneous sequence (row 3, column 1), whereas P-EBM correctly estimates the sequences with simultaneous events (row 3, column 3).

### 2.3. Sequence estimation

For a given dataset, the sequence is estimated using maximum likelihood estimation (Fig, 1.C). This is equivalent to maximum a posteriori probability estimation with uniform prior probabilities on the sequence. When the dataset contains few biomarkers, the sequence can be estimated using exhaustive search, as shown in Fig. 2.

Typically however, datasets with more biomarkers are used and exhaustive search is no longer feasible. In this case, the sequence with maximum likelihood is estimated using stochastic greedy ascent (Young et. al. 2014). The greedy ascent initialises the proposed sequence randomly and iteratively perturbs the sequence, retaining only those with higher likelihood, until convergence, or a maximum number of iterations is reached (Fig. 1C, left).

To maximise the chance of convergence to the global optimum, several initialisations are performed and the sequence with maximum likelihood over all ascents is taken as the estimated sequence. Markov Chain Monte Carlo (MCMC) may also be used to sample additional sequences around the candidate solution (Fonteijn et. al. 2012) and reduce the chance of local maximum (Fig. 1C, right).

Greedy ascent and MCMC utilise a perturbation method to propose different pseudo- random sequences. Below, we describe the perturbation method used for the EBM and its generalisation for the P-EBM. The perturbation techniques for each are summarised in Fig. 3.

**Figure 3.**
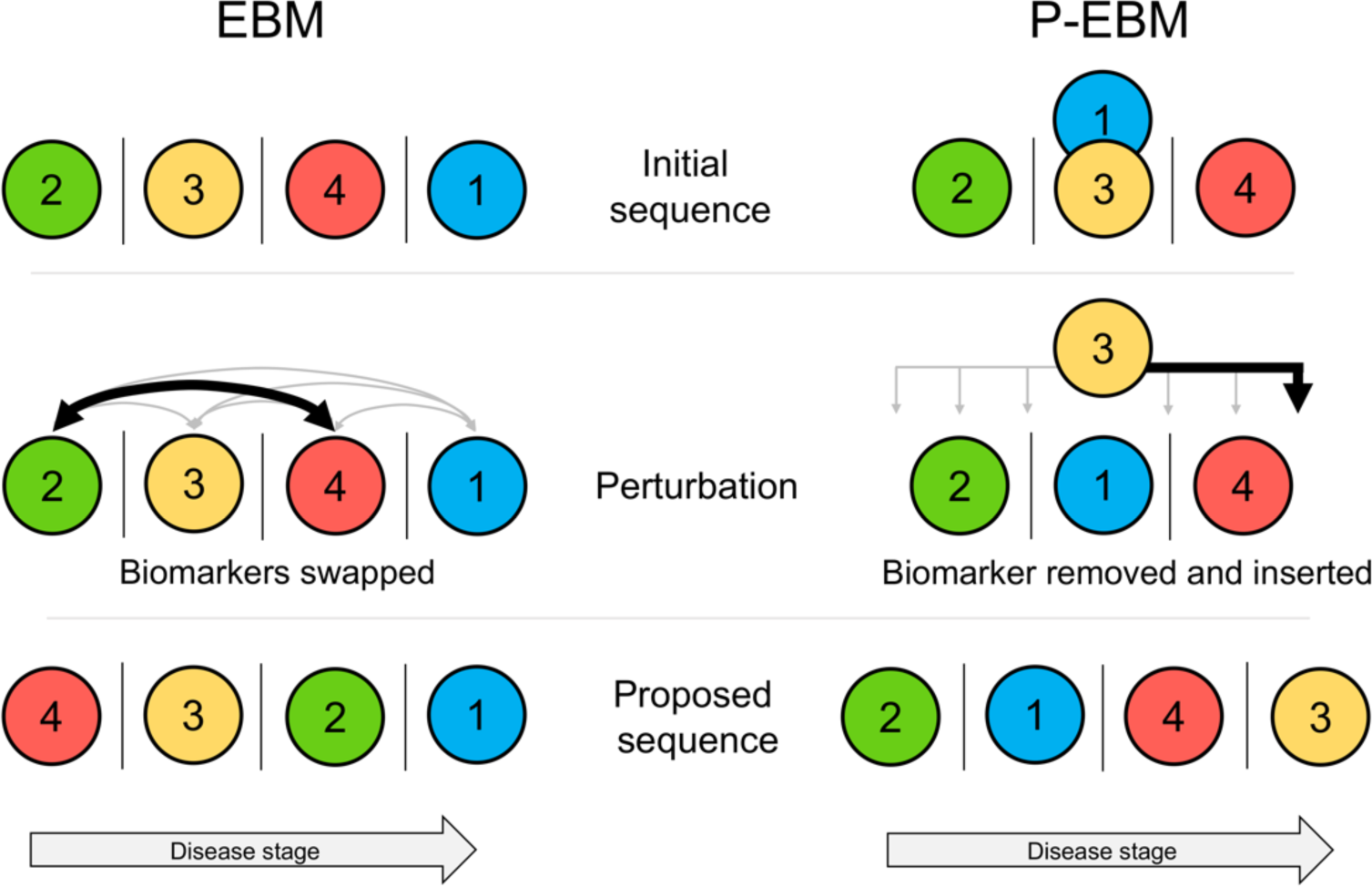
Pictorial overview of the sequence perturbation technique for the EBM and P-EBM. The EBM perturbs an initial sequence by swapping the stages of two biomarker events at random (grey arrows). The P-EBM perturbs an initial sequence by randomly removing one biomarker and replacing it randomly at any of the candidate insertion positions (grey arrows).

#### 2.3.1. Event-based model of disease progression

A proposed sequence is generated by swapping the modelled stages of two biomarker events at random. For example, if the current sequence is (2, 3, 4, 1), then a proposed sequence can be generated by swapping biomarkers 4 and 2, giving the sequence (4, 3, 2, 1) (Fig. 3).

#### 2.3.2. Parsimonious event-based model of disease progression

To enable the list of sets to vary arbitrarily, we update the sequence perturbation method. A biomarker is chosen at random, removed, and is then replaced at a random valid position, such that it generates a valid P-EBM sequence given the arrangement of the other biomarkers. When replaced, it merges with other events at that stage, if there are any. For example, for the sequence ({2}, {1, 3}, {4}), a proposed sequence might be generated by randomly choosing biomarker 3 and replacing it at position 4, giving the sequence ({2}, {1}, {4}, {3}) (Fig. 3). Further example perturbations are shown in Table S1. Note that this perturbation method is compatible with MCMC as it retains the property of symmetric transition probability p(S_0+$_|S_0_) = p(S_0_|S_0+$_).

### 2.4. Positional variance diagram

A positional variance diagram (PVD) is used to report the uncertainty in event positions associated with the maximum likelihood sequences (Fonteijn et. al. 2012).

#### 2.4.1. Event-based model

In the EBM, the PVD utilises samples of the posterior distribution of sequences generated using MCMC. Across MCMC samples, the frequency that a particular biomarker event occurs at a particular stage of the sequence approximates the posterior probability that the biomarker event occurs at that stage. The PVD displays this information in a grid, i.e., for biomarker m and stage k , 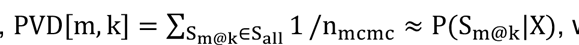, where S_2@-_ denotes a sequence with biomarker m at stage k and S_566_ is the set of sequences generated from MCMC. MCMC samples are generated using the Metropolis Hastings algorithm, with candidate sequences suggested following the perturbation technique described in §2.3.1.

#### 2.4.2. Parsimonious event-based model

The positional uncertainty of P-EBM sequences can be represented in an analogous PVD. Sequences are sampled from the posterior distribution using MCMC, and the frequency over samples for a biomarker event at a particular stage is displayed in the PVD i.e., for biomarker m and stage k , PVD[m, k] ≈ P(S_2@-_|X). To enable the sequence to vary arbitrarily, candidate sequences are suggested following the P-EBM perturbation technique described in §2.3.2. As the sequence can contain simultaneous events, the MCMC can sample sequences with different numbers of stages. As with the EBM, the PVD gives an indication of the uncertainty in the stage at which each biomarker event occurs.

### 2.5. Patient staging

Given an estimated sequence of events, the stage most compatible with a subject’s biomarker measurements can be estimated as the stage with maximum likelihood, as described in (Fonteijn et. al. 2012) and (Young et. al. 2014).

#### 2.5.1. Event-based model

The patient stage is estimated as the stage with maximum likelihood over all stages of the serial sequence estimated by the EBM, 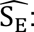

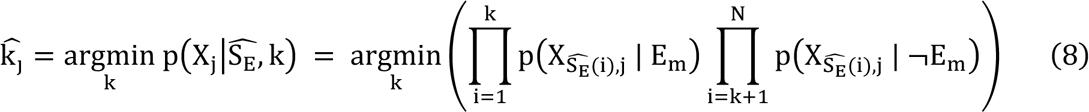

#### 2.5.1. Parsimonious event-based model

The patient stage is estimated from the estimated P-EBM sequence, 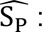

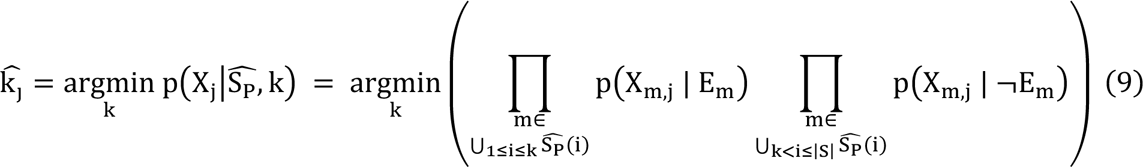

### 2.6. Estimating pre- and post- event state probability density functions

The sequence likelihood requires the pre- and post-event biomarker probability density functions (henceforth referred to as biomarker event pdfs) for each biomarker: 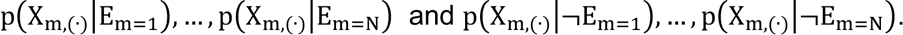. Hypothetically, if each subject’s sequence stage is already known, the event state for each biomarker measurement is also known, and the event pdfs can be fitted to the corresponding subjects’ biomarker measurements. For example, the event pdf for the i’th biomarker, 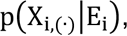, can be fitted to the i’th biomarker measurements from those subjects at a stage equal or greater than the stage that the event occurs for biomarker 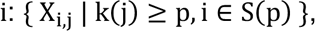, where k(j) is the stage of subject j. However, as the event condition for each biomarker measurement is unknown a priori, then to recover the biomarker event pdfs, a mixture model, p(X_i,(·)_) 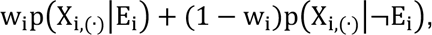, is fit to all subjects’ biomarker measurements, for each biomarker, as in (Fonteijn et. al. 2012, Young et. al. 2014).

## 3. Materials and Methods

### 3.1. Overview

We evaluate the P-EBM and EBM in experiments involving simulated and real data. In simulated data, we firstly evaluate the P-EBM and EBM sequence estimation performance in terms of distance to ground truth and fit quality. We next assess the ability of the EBM PVD’s to disentangle simultaneous events from degree of biomarker inter-subject variation. The capability of P-EBM to stage subjects using less biomarker data, by removing biomarkers whose events are simultaneous and therefore potentially redunandant, is tested by comparing the subject stages estimated on the full dataset to a reduced dataset suggested from the P-EBM estimated sequence. P-EBM is then demonstrated in real data to derive the sequence of biomarker events in sporadic AD and the sequence compared against the EBM.

### 3.2. Simulation experiments

This section describes the simulation process, the set of simulated datasets produced for performance evaluation, and the analysis methods.

#### 3.2.1. Parsimonious event-based forward model

The P-EBM forward model is used to simulate biomarker data as it is more general than the EBM. The required inputs are: (i) the sequence, which may contain serial (single-biomarker) events, simultaneous (multi-biomarker) events, or a mixture of both; (ii) the event pdfs for each biomarker and (iii) the number of subjects in the dataset.

The simulation process proceeds as follows. First, each subject’s stage within the sequence is sampled from a uniform distribution, i.e., Unif{0, |S|}. Then, for each subject j with sequence stage k, biomarker data is generated by sampling from the biomarker event pdfs that correspond to the subject’s sequence stage: abnormal/normal data is sampled if the biomarker event has occurred/has not occurred at that sequence stage i.e. the entry for biomarker m is sampled from p(X_m,(·)_|E_m_) if m ∈ U_1≤i≤k_ S(i), or p(X_m,(·)_,|E_m_) if m ∈ U_k<i≤|S|_ S(i). Sampled biomarker data from N biomarkers and J subjects forms a matrix X, of size N-by-J. All event pdfs are defined as gaussian with a mean of 0 and 1 for pre- and post-event, respectively, and with an equal standard deviation.

#### 3.2.2 Simulation datasets

Using the P-EBM forward model, we simulated datasets with varying numbers and arrangement of simultaneous events and for biomarkers with varying degrees of disease signal, with the aim of comparing sequence estimation performance between the P-EBM and the EBM. To test generalisability of the results, datasets were also simulated with varying numbers of biomarkers and numbers of subjects.

To test performance across a wide spectrum of sequence types, data was generated from ground truth sequences with differing proportions of simultaneous events: 0, 0.25, 0.5, 0.75 and 1. Simultaneous events were either assigned to a single stage or multiple stages. For the single-stage scenario, all simultaneous events were assigned to the first disease stage. For the multi-stage scenario, each stage with simultaneous events contained a quarter of the biomarkers in the full dataset. Datasets with different degree of disease signals were generated by setting the biomarker inter-subject standard deviation at 0.25, 0.5, 0.75 and 1 (disease effect sizes of 4, 2, 1.33 and 1, respectively).

To test whether the trends of performance differences were consistent in datasets of different sizes, we also simulated datasets which varied the number of biomarkers and number of subjects. The number of biomarkers was varied across the range typically used for EBM studies (Fonteijn et. al. 2012, Young et. al. 2014, Venkatraghavan et. al. 2017, Young et. al. 2018, Oxtoby et. al. 2018, Firth et. al. 2020, Gabel et. al. 2020): 12, 24 and 36; and was evenly divisible by the proportion of simultaneous events. The number of subjects was varied to cover realistic variations in the size of neuroimaging data collections (175, 350, 750).

For each combination of simulation settings, 100 datasets were generated, and both the P-EBM and the EBM were applied to estimate the maximum likelihood sequence and corresponding PVD. Subjects were then staged within the P-EBM sequence using the full dataset and a reduced dataset suggested by removing biomarkers whose events are simultaneous.

#### 3.2.3. Sequence estimation

The number of initialisations and iterations of the greedy ascent were increased for datasets of more biomarkers to account for the increasing size of the sequence search space and maximise the chance of identifying the global maximum: 40 initialisations and 3000 iterations were used for datasets with 12 biomarkers; 50 initialisations and 4000 iterations were used for datasets with 24 biomarkers, and 60 initialisations and 5000 iterations were used for datasets with 36 biomarkers. To further improve the chance of finding the global maximum likelihood sequence, an additional 10^6^ MCMC iterations were performed starting from the sequence with highest likelihood identified by greedy ascent. The sequence with maximum likelihood across MCMC samples was then taken as the estimated sequence. Event pdfs were estimated using a gaussian mixture model, as described in (Young et. al. 2014).

#### 3.2.4. Positional uncertainty

For each simulated dataset, and following sequence estimation, P-EBM and EBM PVDs were calculated from an additional 10^6^ MCMC samples, as described in §2.4.

#### 3.2.5. P-EBM staging with reduced data

To stage subjects with a reduced dataset using P-EBM, a reduced sequence and associated dataset is derived from the estimated sequence and its associated dataset. Firstly, qxw reduced sequence was defined by randomly selecting one biomarker event for each stage of the full sequence and discarding all others. Subsequently, the reduced dataset was defined by retaining only the biomarker data for the events in the reduced sequence. Subjects were then staged within the full sequence using the full dataset, and within the reduced sequence using the reduced dataset, by maximising the subjects stage likelihood according to Eq. 9.

#### 3.2.6. Analysis of sequence estimation performance

We evaluate sequence estimation performance using two metrics. The Kendall tau distance between the estimated sequence and the ground truth sequence was used to evaluate the (dis)-similarity of estimated sequences. Secondly, fit quality was evaluated using the log of the ratio of likelihoods (LLR) between the P-EBM estimated sequence and the EBM estimated sequence.

##### 3.2.6.1. Kendall Tau Distance

The Kendall tau distance has been previously to assess EBM sequence estimation performance (Huang & Alexander 2012, Young et. al. 2015, Venkatraghavan et. al. 2017, Khatam et. al. 2022, Young et. al. 2023, Tandon et. al. 2023). The metric quantifies how far an estimated sequence is from the ground truth sequence in terms of pair-wise ranking differences among all pairs of biomarker events.

We adopt a version of the Kendall tau distance suitable for both serial sequences and sequences that contain simultaneous events (Fagin et. al. 2008). The distance was defined as the fraction of pair-wise biomarker event with a different order over all pairs of events, with the ordering of two biomarker events, *A* and *B*, represented by an integer (1: *A* before *B*, 2: *A* after *B*, 3: *A* and *B* are simultaneous). A disagreement is observed if a pair of events have a different order. This formulation is the Kendal tau distance for partial rankings with a penalty term of 1 (Fagin et. al. (2008).

Lower distance means the estimated sequence has higher similarity to the ground truth sequence. Distance ranges from 0 (correct sequence; no pair-wise event orders disagree) to 1 (incorrect sequence; all pair-wise event orders disagree). The Kendall tau distance was calculated for each estimated sequence.

As the original EBM cannot estimate simultaneous events, the minimum distance the EBM sequence can achieve is limited by the number of simultaneous events in the ground truth sequence. For each stage of the ground truth sequence that contains simultaneous events, all possible within-stage pair-wise event orders will differ from the EBM estimated sequence, while all other pair-wise event orders can match. Given the ground truth sequence, S_9:_, and the set of its stages that contain simultaneous events, m = {j| |S_9:_(j)| > 1}, the lower bound on the distance of an EBM estimated sequence is:

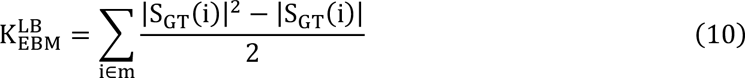

From Eq. 10, the original EBM can only achieve the minimum distance of 0 for serial ground truth sequences. For sequences containing simultaneous events, the minimum distance for EBM estimated sequences increases as the number of simultaneous events per stage and the number of stages containing simultaneous events increases.

As the P-EBM can estimate sequences with an arbitrary arrangement of simultaneous events, the minimum distance of P-EBM estimated sequences is 0.

##### 3.2.6.2. Fit quality

The LLR between the P-EBM estimated sequence and the EBM estimated sequence were calculated as ln [ p+X|Ŝ_P-EBM_/p(X|Ŝ_EBM_)]. LLR>0, or <0, indicates superior fit of the P- EBM, or EBM model, respectively. The LLR value was calculated for each simulated dataset on which the P-EBM and EBM sequences were estimated.

#### 3.2.7. Analysis of EBM positional uncertainty

We compare the extent to which the uncertainty of EBM PVDs are influenced by simultaneous events and biomarker inter-subject variation. PVD uncertainty was quantified as the mean positional standard deviation across the PVD matrix and plotted for varying proportions of simultaneous events and varying biomarker inter-subject standard deviation.

#### 3.2.8. Analysis of staging with reduced data

For each simulated dataset, the Pearson correlation across subjects between the stage estimated on the full and reduced dataset, and the percentage of correctly staged subjects, was calculated.

### 3.3. Application to sporadic Alzheimer’s disease

We tested the ability of the P-EBM to estimate a plausible sequence of biomarker abnormalities in sporadic AD and compared the sequence and quality of fit to the EBM. We also assessed the capability for P-EBM to suggest a reduced biomarker subset for patient staging.

#### 3.3.1. ADNI data source

We used a sporadic AD dataset derived from the Alzheimer’s disease Neuroimaging Initiative (ADNI) and described in (Young et. al. 2014). Data was obtained from the ADNI database (adni.loni.usc.edu) using LONI (www.loni.ucla.edu/ADNI/), in compliance with the ADNI data use agreement. Written informed consent was given for all participants, and the study was approved by the Institutional Review Board at each participating institution, in accordance with their ethical standards and the 1964 Helsinki declaration.

The ADNI was launched in 2003 as a public-private partnership with the goal to test whether serial magnetic resonance imaging, positron emission tomography, other biological markers, and clinical and neuropsychological assessment can be combined to measure the progression of mild cognitive impairment and early Alzheimer’s disease. For up-to-date information, see http://www.adni-info.org. ADNI data captures a wide range of demographic diversity due to ADNI’s study design, which maximises participation of under-represented minorities.

#### 3.3.2. Subjects and biomarkers

The dataset is identical to that used to demonstrate application of the EBM to sporadic AD (Young et. al. 2014). It contains 325 subjects with purely cross-sectional data for 12 biomarkers, consisting of 83 patients with AD-dementia, 218 subjects with mild cognitive impairment and 54 cognitively normal controls (demographics in Table S2). Subjects were selected according to those who underwent CSF examination at baseline (tau, phosphorylated tau and amyloid-β1-42), standardized cognitive assessment at baseline (Mini-Mental State Examination (MMSE) (McKhann et. al. 1984), the Alzheimer’s Disease Assessment Scale – Cognitive Subscale (ADAS-Cog) (Rosen et. al. 1984) (modified 13- item ADAS-Cog, which omits Item 13), and the Rey Auditory Verbal Learning Test (Rey, 1958) (immediate recall score, i.e., the sum of trials 1 to 5)), and 1.5 T MRI imaging at baseline that passed quality control (regional volumes of entorhinal cortex, hippocampus, mid-temporal gyrus, fusiform gyrus, ventricles and whole-brain).

#### 3.3.3. Biomarker processing

Data acquisition and processing of ADNI biomarkers has been described on the ADNI website (adni.loni.usc.edu) and in previous reports (Jack Jr. et. al. 2008, Shaw et. al. 2009, Crane et. al. 2012). Brain volumes were calculated using FreeSurfer Version 4.3 (http://surfer.nmr. mgh.harvard.edu/). For EBM event pdf estimation, cerebrospinal fluid (CSF) total tau and phosphorylated tau were log-transformed to improve normality (Young et. al. 2014). Regional volumes were averaged over hemispheres and normalised by intracranial volume to control for individual differences in head size.

#### 3.3.4. Biomarker event pdf estimation

For each biomarker, a constrained gaussian mixture model was fit to the cognitively normal and AD patients’ measurements, as in (Young et. al. 2014). The constraints ensure a robust fit in the case where the distributions of cognitively normal and patient populations overlap significantly. The standard deviations of the pre-event and post-event pdfs are constrained to be less than or equal to that of the cognitively normal or AD group, respectively, and the means of the pre-event and post-event pdfs are constrained to be less extreme (normal/abnormal) than the cognitively normal or AD groups, respectively.

#### 3.3.5. Sequence estimation

The maximum likelihood P-EBM and EBM sequences were estimated using greedy ascent and MCMC. Greedy ascent was used to find an initial estimate of the sequence using 40 initialisations each with 3000 iterations. MCMC with 10^6^ samples was then used to refine the estimate.

#### 3.3.6. Positional variance

To estimate positional uncertainty, an additional 10^6^ MCMC sequence samples were generated. The P-EBM and C-EBM PVD’s were calculated as described in §2.4.

#### 3.3.7. Subject staging with reduced data

ADNI subjects were staged using the full dataset, and a reduced dataset produced as described in §3.2.5 and according to Eq. 9. The Pearson correlation and percentage of correctly staged subjects are reported.

## 4. Results

### 4.1. Simulation experiments

#### 4.1.1. Kendall tau distance

Fig. 4 shows the Kendall tau distance of estimated sequences against the ground truth over a range of number of simultaneous events and degree of biomarker inter-subject standard deviation. We show results here for datasets of 12 biomarkers and 350 subjects, which is representative of the AD dataset (§4.2). The equivalent plots across all dataset sizes are shown in Fig. S1 and Fig. S2.

**Figure 4.**
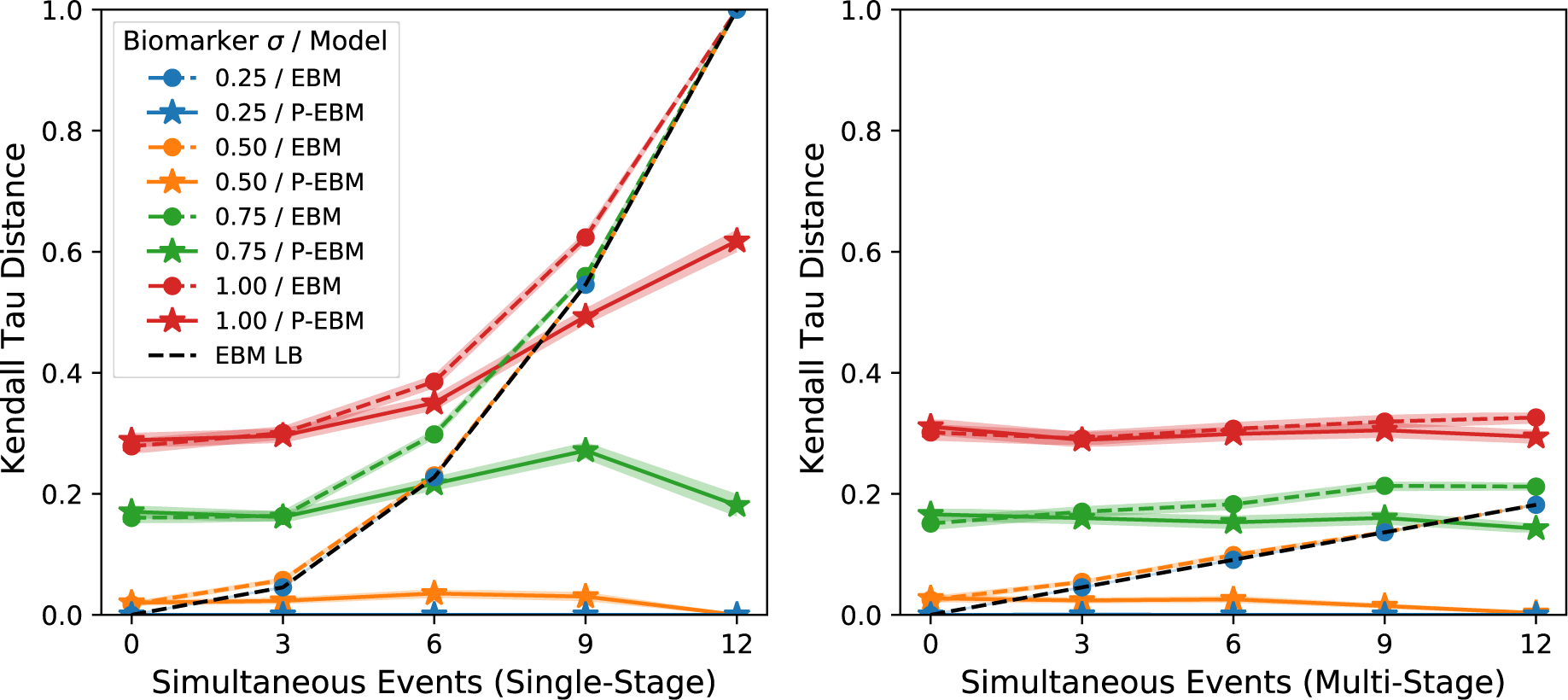
Kendall tau distance of the estimated sequences against the ground truth sequence for simulated datasets of 12 biomarkers and 350 subjects. **Left:** Ground truth sequences generated containing a single stage with varying number of simultaneous events. **Right**: Ground truth sequences generated containing multiple stages with 3 simultaneous events each. Lines and shaded regions show the mean and standard error across the 100 simulated datasets. The dotted line shows the lower bound on the EBM distance metric.

For P-EBM, under almost all sequence scenarios, the majority of pairwise event ordering categories (before, after, or simultaneous) were correctly assigned. For serial sequences (number of simultaneous events=0), both the P-EBM and EBM performed equally, with both methods achieving low Kendall tau distance. However, for sequences with a single stage of simultaneous events (Fig. 4, left) or varying number of stages with simultaneous events (Fig. 4, right), the Kendall tau distance of P-EBM-estimated sequences was lower than or equal to the EBM.

For datasets with low inter-subject biomarker standard deviation (≤0.5, corresponding to disease effect size of ≥2), the P-EBM almost perfectly estimated the ground truth sequence, as shown a distance equal to close to 0. However, the EBM shows a systematic increase in distance with the number of simultaneous events:- the lower bound on the EBM distance increased as the number of simultaneous events increased, and did so more rapidly when simultaneous events were assigned to a single stage rather than multiple stages. Despite the EBM’s inability to estimate sequences containing simultaneous events, it did achieve the lower bound on distance when inter-subject standard deviation was low, and showed reasonable distance to the ground truth when the number of simultaneous events was small.

For both models, distance tended to increase as biomarker inter-subject standard deviation increased (disease effect size decreased). Trends of differences were similar for datasets of different number of biomarkers and number of subjects, with reduced distance observed as the number of subjects increased (Fig. S1, S2).

#### 4.1.2. Fit quality

Fig. 5 shows the LLR between P-EBM and EBM estimated sequences as the number of simultaneous events increased. We report results here for datasets of 12 biomarkers and 350 subjects, with plots across all dataset sizes shown in Fig. S2.

**Figure 5.**
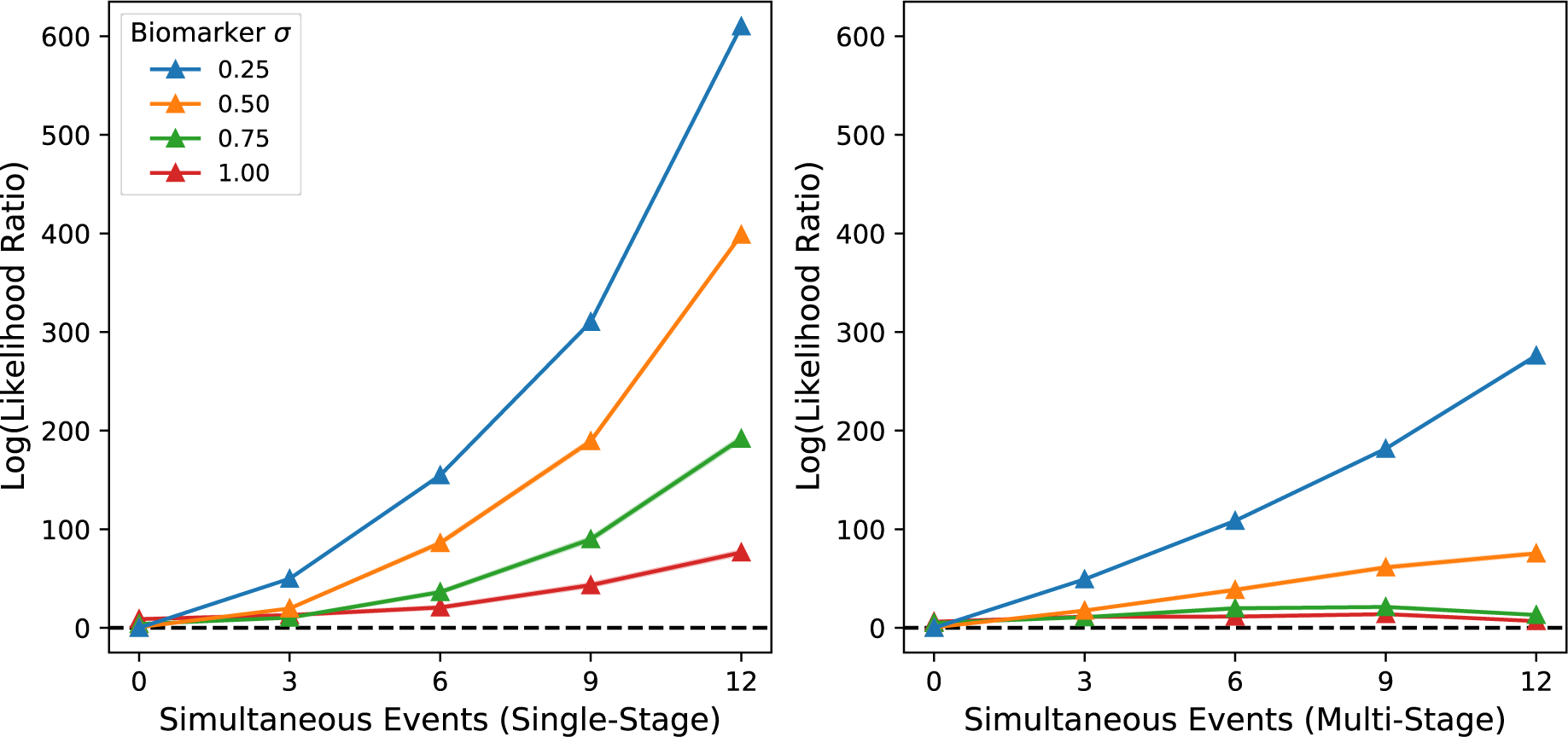
Log of the likelihood ratio between the P-EBM estimated sequence and the EBM estimated sequence for simulated datasets with 12 biomarkers and 350 subjects. **Left:** Ground truth sequences generated containing a single stage with varying number of simultaneous events. **Right**: Ground truth sequences generated containing multiple stages with 3 simultaneous events. Lines and shaded regions show the mean and standard error across the 100 simulated datasets. LLR>0 indicates superior fit quality of the P-EBM compared to the EBM.

**Figure 6.**
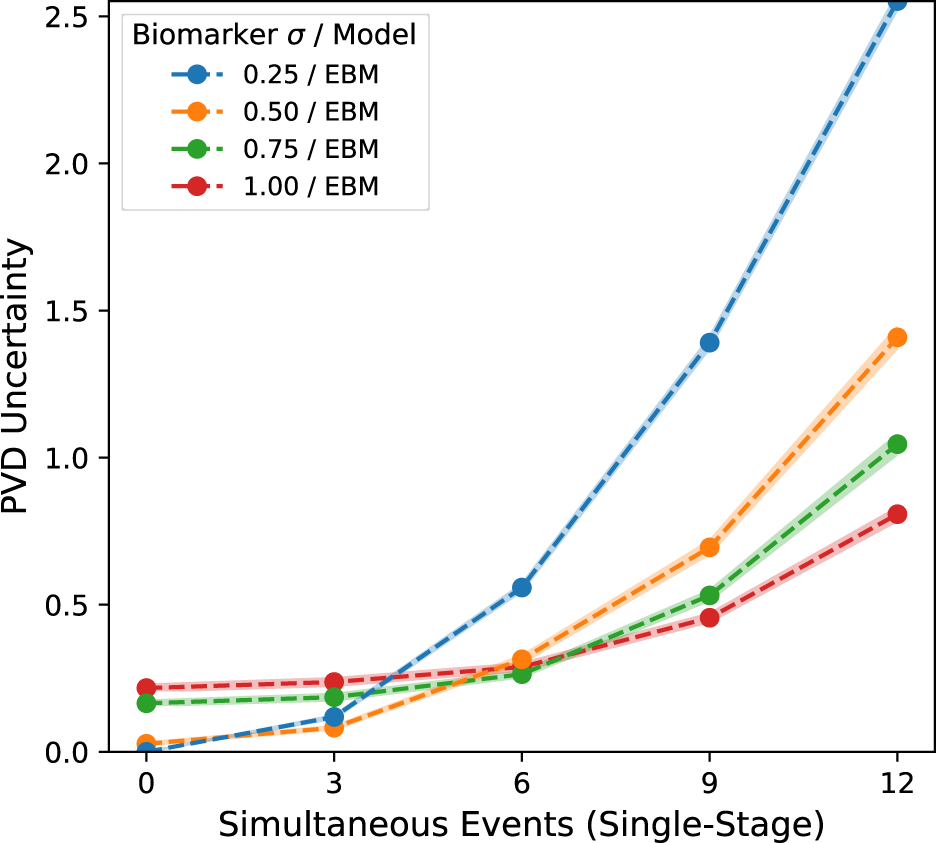
EBM PVD uncertainty (mean positional standard deviation) of estimated sequences for simulated datasets with 12 biomarkers and 350 subjects. **Left:** Ground truth sequences generated containing a single stage with varying number of simultaneous events. **Right**: Ground truth sequences generated containing multiple stages with 3 simultaneous events. Lines and shaded regions show the mean and standard error across the 100 simulated datasets.

The LLR was above 0 and increased as the number of simultaneous events increased, demonstrating that the P-EBM estimated sequences provide an increasingly closer fit to the data than the EBM as the number of simultaneous events increases. The LLR increases more rapidly with the number of simultaneous events for the single-stage scenario as compared to the multi-stage scenario.

The LLR tended to decrease as the biomarker inter-subject standard deviation increased, indicating that for noisy data instances the relative difference in fit quality is reduced. Similar trends were observed in datasets with varying numbers of biomarkers and numbers of subjects (Fig. S3, S4).

#### 4.1.4. EBM positional uncertainty

The EBM PVD uncertainty depended strongly on the number of single stage simultaneous events, whereas it showed less dependence on the number of multi-stage simultaneous events. Therefore, we report the single stage scenario here, with the dependence on multi-stage events shown in Fig. S6.

As the number of singe stage simultaneous events increases, the degree of PVD uncertainty also increases. However, the PVD uncertainty also shows a dependence on the biomarker inter-subject standard deviation. The PVD uncertainty furthermore depended on the number of stages with simultaneous events to a lesser degree. Similar dependence patterns are observed in datasets with varying numbers of biomarkers and numbers of subjects (Fig. S5, S6).

#### 4.1.5. P-EBM staging with reduced data

Fig. 7 shows the percentage of subjects assigned to the correct stage when using the reduced dataset for the case of 12 biomarkers and 350 subjects. The percentage of correctly estimated subject stages in the reduced dataset was high (>70%) for all sequences and biomarker inter-subject standard deviations, decreasing by 5-10% as the number of simultaneous events increased to its maximum. Similar trends were observed in datasets of different sizes (Fig. S7, S8).

**Figure 7.**
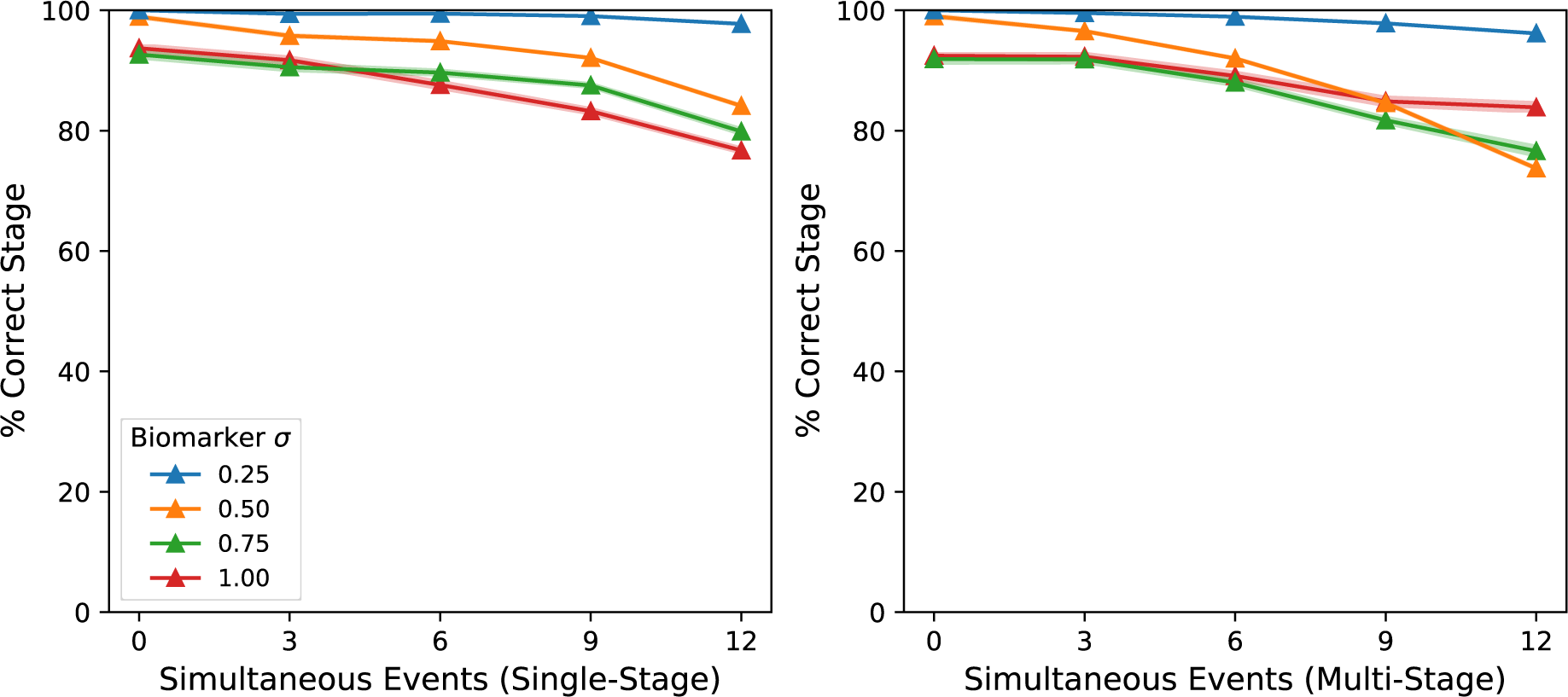
Percentage of subjects assigned to the correct stage of the P-EBM sequence using the reduced dataset for sequences of 12 biomarkers and 350 subjects. **Left:** Ground truth sequences generated containing a single stage with varying number of simultaneous events. **Right**: Ground truth sequences generated containing multiple stages with 3 simultaneous events. Lines and shaded regions show the mean and standard error across the 100 simulated datasets. LLR>0 indicates superior fit quality of the P-EBM compared to the EBM.

Those subjects that assigned to an incorrect stage in the reduced dataset were still assigned to a similar stage as in the full dataset, as indicated by the high Pearson correlation coefficient between the stage for the full and reduced datasets (Fig. S9, S10) and the high number of entries in the diagonal of the joint histogram (Fig. 8).

**Figure 8.**
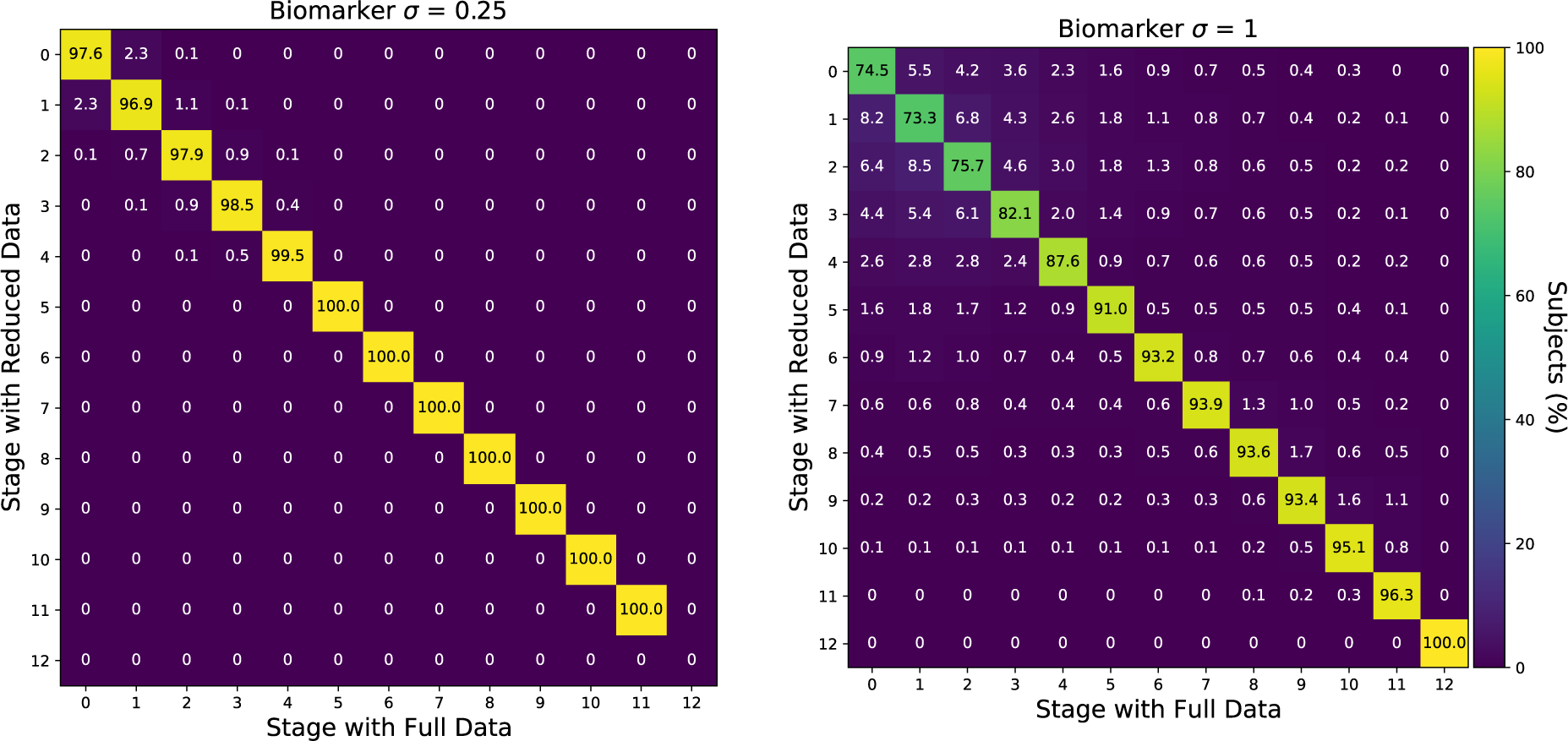
Normalised joint histogram of the estimated subject stage in the reduced and full dataset for simulated datasets of 12 biomarkers and 350 subjects, accumulated across all single and multi-stage sequences containing simultaneous events. Each square shows the % of patients assigned to each stage for those patients assigned to the stage indicated by the column when using the full dataset.

The reduced dataset selects data for one biomarker from each P-EBM sequence stage.

The early stages were sometimes mis-assigned when biomarker variation was high (inter-subject standard deviation=1, corresponding to disease signal=1; Fig. 8, right). These stages correspond to shorter sequences where the number of simultaneous events is highest and the reduction in the number of biomarkers is greatest.

### 4.2. Application to Alzheimer’s disease

#### 4.2.1. Sequence of AD progression

To aid comparison to simulation results, effect sizes and the corresponding simulated biomarker inter-subject standard deviation for each biomarker are reported in Table S3. Effect sizes ranged from 1.2 to 3.4, corresponding to simulated biomarker inter-subject standard deviations of 0.8 to 0.3.

Fig. 9 shows the sequence of AD progression and its uncertainty, as estimated by the P-EBM, and the EBM. As expected, the EBM estimated a serial sequence with a single biomarker event at each disease stage (log-likelihood = 3620). The sequence matches that reported in the original application of the EBM to sporadic AD (Young et. al. 2016 Supplementary).

**Figure 9.**
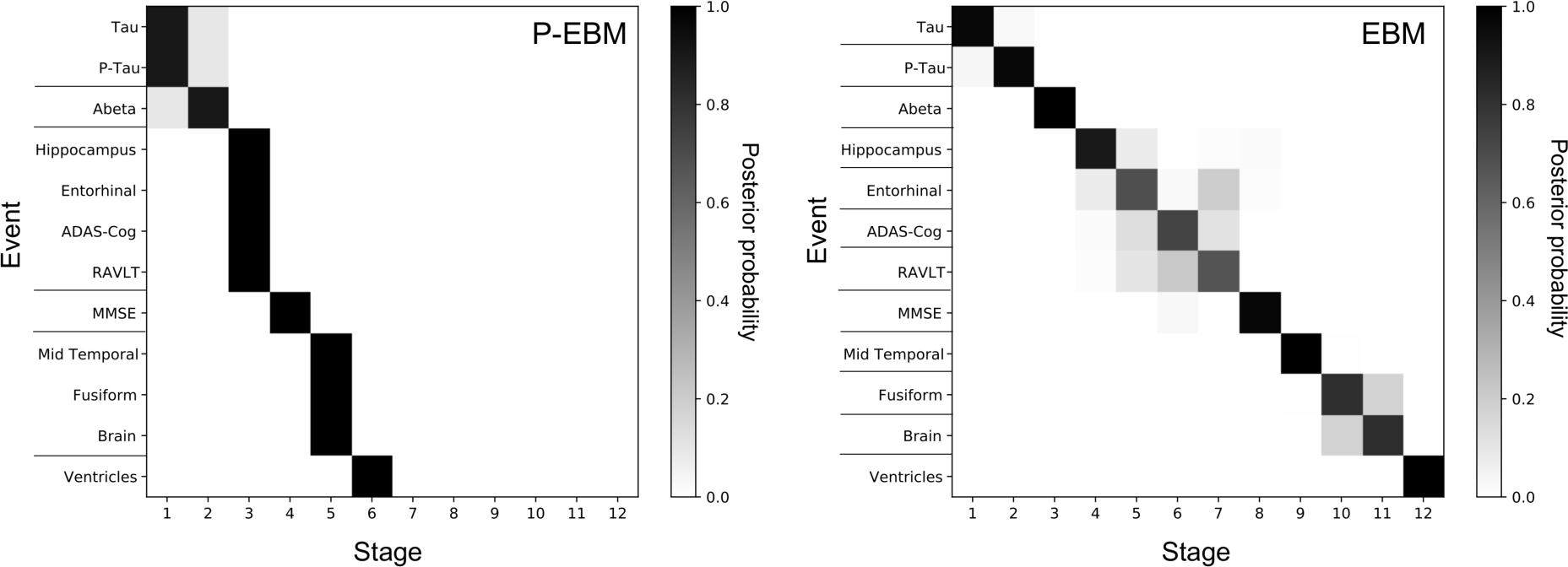
The estimated sequence of biomarker events in sporadic AD and its uncertainty for the P- EBM and EBM. The estimated sequence is shown by the PVD row ordering, with lines separating the events in each stage. PVD entries show the posterior probability of each biomarker event (row index) at each stage (column index).

The P-EBM estimated a sequence with higher likelihood than the EBM (log-likelihood = 3691), consisting of both serial and simultaneous events. The P-EBM sequence begins with simultaneously abnormal CSF tau and phosphorylated tau (stage 1), followed by abnormal CSF amyloid-β1-42 (stage 2), followed by simultaneously abnormal hippocampal volume, entorhinal cortex volume, ADAS-Cog scores, and RAVLT scores (stage 3). MMSE then becomes abnormal (stage 4), followed by simultaneous events for mid-temporal gyrus volume, fusiform gyrus volume and whole brain volume (stage 5). Ventricle volume is the last event in the sequence (stage 6).

The EBM PVD suggests different degrees of uncertainty in the serial ordering of biomarkers. For biomarkers with relatively higher effect sizes, this uncertainty may suggest but not confirm the presence of simultaneous events. This is the case for the event stages of entorhinal cortex volume, RAVLT and ADAS-Cog, whose effect sizes are among the highest (2.0, 3.2, 2.7, corresponding to biomarker standard deviations of 0.5, 0.31, 0.37). However, for biomarkers with relatively low effect sizes, such as fusiform gyrus and brain volume (1.2 and 1.7, corresponding to biomarker standard deviations of 0.83 and 0.58 in the simulations), the EBM PVD cannot disentangle the presence of simultaneous events from uncertain serial events for these biomarkers.

The P-EBM PVD suggests high confidence in the event orderings of most biomarkers, with almost all MCMC samples being the maximum likelihood sequence, which shows that the estimated sequence has a much higher likelihood than the other sequences sampled during the MCMC step. CSF total tau, phosphorylated tau and amyloid-β1-42 events are less certain, with a small proportion of positional variance between events.

#### 4.2.2. Bootstrap resampling

The sensitivity of the P-EBM and EBM sequence to the initial dataset was assessed using bootstrap resampling. The maximum likelihood sequence was estimated for 100 bootstraps by re-sampling the original data with replacement. Fig. 10 shows the variation in event stages over the estimated sequences. The PVD shows high correspondence to the original sequence for both P-EBM and EBM, with the exception that the maximum likelihood sequence estimated in the resampled data often staged worsening performance on ADAS-Cog and RAVLT tests prior to volumetric loss of hippocampus and entorhinal cortex.

**Figure 10.**
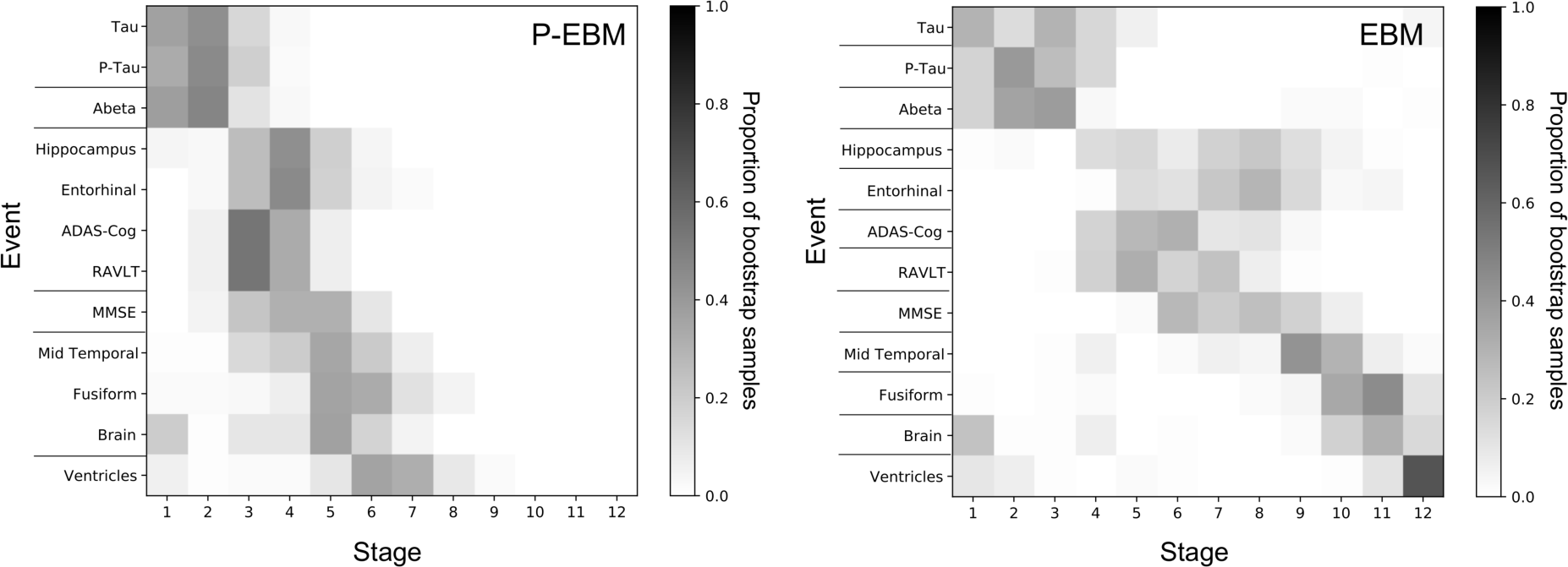
Positional variance diagrams from bootstrap resampling, showing the distribution of event stages over sequences estimated from 100 bootstrap samples of the original data. Each entry shows the proportion of times the event appeared at a particular stage.

#### 4.2.3. Staging subjects with the reduced dataset

The reduced dataset selected one biomarker from each P-EBM sequence stage randomly, resulting in six biomarkers:- P-Tau, Abeta, ADAS-Cog, MMSE, Brain volume and Ventricle volume; a 2-fold decrease in the number of biomarkers. Subject stages estimated on the reduced dataset closely match the stages estimated on the full dataset (Fig. 11), with the Pearson correlation coefficient being 0.94 (*p-value* = 6.7x10^-156^) and 88.3% of subject stages matching overall.

**Figure 11.**
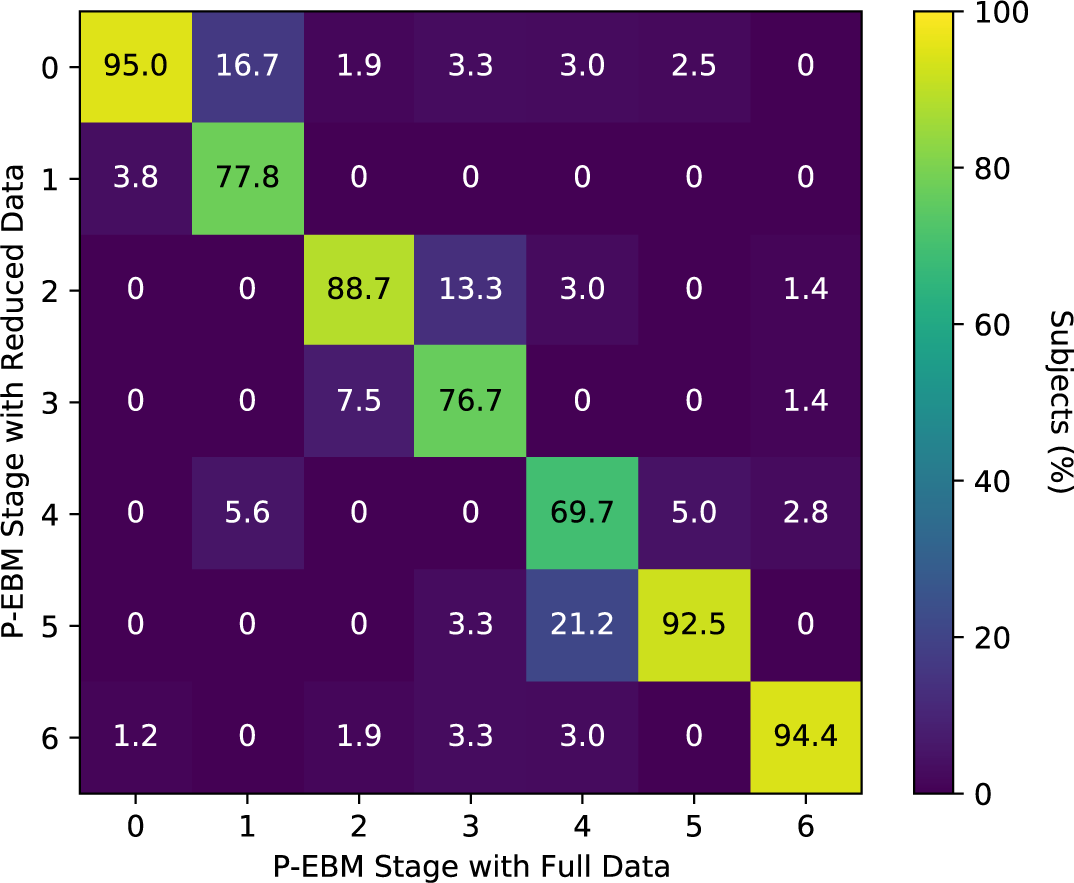
Normalised joint histogram of the estimated subject stage in the reduced and full ADNI dataset.

## 5. Discussion

### 5.1. Summary

We introduce the P-EBM, a generalisation of the event-based model that enables the estimation of shorter, more parsimonious disease progression sequences that contain fewer stages than the number of input biomarkers. Simulation analysis demonstrates high sequence estimation performance with varying numbers of simultaneous events and under a range of realistic experimental scenarios. Staging analysis shows that P-EBM can suggest a reduced data subset to accurately stage subjects within a shorter sequence, achieving comparable performance to the full dataset. In sporadic AD, the P-EBM reconstructs a plausible sequence containing simultaneous events which closely explains the biomarker measures. The following paragraphs interpret the simulation analysis, discuss the P-EBM sequence inferred in sporadic AD, compare the P-EBM to other event-based models and highlight future challenges and opportunities.

### 5.2. Simulation analysis

Simulation analysis shows that the P-EBM has high fidelity for estimating sequences of serial events and sequences containing simultaneous events (Fig. 4, S1-4). P-EBM estimated sequences had low distance to ground truth and higher likelihood than EBM estimated sequences. The P-EBM’s ability to reconstruct a more parsimonious disease progression sequence widens the applicability of the EBM to larger datasets, with potential for improving disease understanding and generating practical staging systems.

The P-EBM’s broader capability is afforded by its large sequence search space, which we show is practically feasible to traverse and optimise. While the EBM search space rises factorially as N!, with N being the number of biomarkers in the input dataset, the P-EBM search space rises super-factorial as ∑* k! S(N, k), where S is Sterling’s number of the second kind (Fishburn 1985, Fagin et. al. 2006, OEIS 2022). The significant computational resources required to optimise the likelihood over the relatively large sequence search space shows little drawback in practice:- P-EBM sequences reached the lower bound on the Kendall tau distance, suggesting convergence to the maximum likelihood, and the optimisation was observed in reasonable time, with the sequence estimation and PVD derivation together requiring less than 5 minutes per dataset.

The P-EBM and the EBM showed differential performance changes as the number of simultaneous events increased. EBM sequence estimation systematically worsened as the number of simultaneous events increased. This is because the lower bound on the achievable distance of EBM estimated sequences increases with the number of simultaneous events (§3.2.6.1 and Eq. 10). In contrast, the lower bound on the achievable distance of P-EBM sequences is 0. In the presence of simultaneous events, P-EBM successfully estimated a range of sequence types, including those varying numbers of stages containing simultaneous events and varying number of events within those stages, and with near-perfect accuracy when biomarker variance was low (Fig. 4, S1, S2).

Interestingly, P-EBM performance tended to increase for datasets produced with more simultaneous events (Fig. 4, S1, S2). This is explained by the increase in the number of subjects per disease stage. This accentuates the contrast in sequence likelihood between the true generative sequence and other proposal sequences, effectively reducing the chance that the data sample appears like a different sequence to its ground truth. As a consequence, P-EBM is well suited for biomarker datasets where the expected number of simultaneous events is high.

PVD simulation analysis shows that the EBM can suggest, but not confirm, the presence of simultaneous events. Indeed, this suggestion motivated our development of P- EBM. Both higher biomarker variance and more simultaneous events affected the EBM PVD uncertainty (Fig. 6). Analysing the EBM PVDs together with disease effect sizes may be more useful for suggesting the presence of simultaneous events than analysing the PVD by itself. This inference may however be complicated by variation of disease effects across biomarkers, a feature of real datasets which was not analysed in this study. In contrast, P- EBM PVDs maintained high positional certainty across the full range of experiments, almost independent of the proportion of simultaneous events and biomarker variance (Fig. 6). This shows that the P-EBM PVD can indicate high confidence in sequences containing simultaneous events, which would otherwise appear as more ambiguous serial sequences when examining the EBM PVD.

The P-EBM’s ability to infer simultaneous events may be applied to suggest biomarker subsets for patient staging, with the rationale that if two or more events occur simultaneously, then only one biomarker within the group is needed to stage a subject within the sequence. We tested this idea by applying a simple heuristic to identify the biomarker subset from a larger dataset using the P-EBM sequence. Subject staging using this reduced dataset is highly accurate, with reductions in the number of required biomarkers up to a factor of N showing high Pearson correlation and accuracy compared to the full dataset (Fig. 7, 8, 11, S7, S8). This demonstrates a potential for P-EBM to suggest efficient biomarker collection strategies for patient staging, with fewer biomarkers reducing the time and cost of data collection, which may be useful in clinical applications. Future work will evaluate the minimal number of biomarkers required for acceptable diagnostic and prognostic purposes.

### 5.3. Sporadic AD progression

P-EBM is designed to estimate the useful stages of disease progression independently from the input dataset. In sporadic AD, P-EBM estimated a shorter, more parsimonious sequence comprising 6 stages of biomarker progression, whereas the EBM is forced by design to identify 12 stages (the number of input biomarkers). Despite its lower complexity, the P-EBM sequence provides a superior fit to the data than the EBM sequence, as illustrated by the large difference in their log-likelihoods (§4.2.1). Indeed, simulation analysis show high performance of P-EBM for datasets of this size.

In sporadic AD, P-EBM estimated a plausible sequence of biomarker events which broadly agrees with recognised pathological changes. The main sporadic AD subtype is thought to begin with abnormal accumulation of amyloid-β and tau proteins in the hippocampus and entorhinal cortex. These regions later undergo atrophy (neuronal loss and macroscopic volume reductions), leading to cognitive decline. Brain atrophy later occurs in other temporal lobe regions and then more widely throughout the brain (Jack Jr. et. al. 2010, Jack Jr. et. al. 2013). Taken together, the fit quality indicators, comparisons to simulated data and the broad agreement with established pathological changes, suggests that the P- EBM has the potential in practice to estimate parsimonious sequences and identify latent stages of disease from relatively high dimensional datasets. Below, the biomarker profiles of each P-EBM stage are analysed in more detail.

AD is characterised by abnormal accumulation of amyloid plaques and neurofibrillary tangles of tau protein in the brain (Braak and Braak 1991), processes which lead to decreased CSF concentration of amyloid-β1-42 and increased CSF concentration of total tau and phosphorylated tau. The P-EBM suggests that abnormal CSF tau and phosphorylated tau occur simultaneously. This is consistent with previous EBM studies reporting high uncertainty in their serial ordering (Young et. al. 2014, Khatami et. al. 2022) and minimal latency differences (Wijeratne et. al. 2023). Remarkably, both the P-EBM and EBM staged decreased CSF total tau and phosphorylated tau prior to decreased CSF amyloid-β1-42, despite evidence suggesting abnormal CSF amyloid-β1-42 concentration precedes abnormal tau concentrations (Bateman et. al. 2012). As reported in (Young et. al. 2014), this could be explained by ADNI’s inclusion of individuals who are negative for apolipoprotein-E (APOE) genetic risk factor as well as amyloid-negative individuals. The sample therefore includes those undergoing normal aging, in addition to those with potentially presymptomatic non-AD neurodegeneration such as dementia with Lewy bodies or frontotemporal dementia, where early tau deposition is observed (Braak & Del Tredici 2011, Kok et. al. 2009).

The P-EBM suggests that decreased hippocampal and entorhinal cortex volumes and worsening performance on ADAS-Cog and RAVLT are better represented as simultaneous rather than serial events. This is plausible given the high temporal correlation between regional volumetric loss and cognitive test scores (Jack Jr. et. al. 2013) and the high sensitivity of these tests to cognitive decline. The simultaneity of these events is also consistent with EBM studies reporting high uncertainty among the event stages of these biomarkers (Young et. al. 2014) and small latency differences (Wijeratne et. al. 2023). As volumetric reductions reflect cumulative neuronal loss, it may be expected that alternative biomarkers such as blood concentration of N-acetyl-aspartate (Xu et. al. 2008) or diffusion imaging measures of neuron density (Alexander et. al. 2019) may identify earlier stages of neurodegeneration. Future P-EBM work will investigate this. Detecting subtle neurodegeneration at the precipice of cognitive decline could identify individuals to enrich clinical trials of secondary preventative interventions (Andrieu et. al. 2013) that aim to slow or prevent further neuronal loss prior to irreversible brain damage.

The reduced biomarker subset identified by the P-EBM sequence staged subjects within the sequence with high consistency to the full dataset (Fig. 11). Using a single biomarker event at each P-EBM stage, subjects were staged using only six biomarkers, compared to the twelve that otherwise required for the full dataset. This demonstrates the practical capability of P-EBM for staging subjects using less data, which may be useful in resource-scarce applications, such as in clinical settings.

### 5.4. Comparison to other event-based models

P-EBM offers several conceptual advantages over other EBM models. The probabilistic events cascade model (Huang & Alexander 2012) and discriminative EBM (Venkatraghavan et. al. 2017) both inherit the serial events assumption from the EBM. Temporal EBM (Wijeratne et. al. 2022) estimates the variable time intervals between stages and their uncertainty, implicitly handling overlapping events, but requires short-term longitudinal data which is not always available. Scaling EBM (Tandon et. al. 2023) performs feature selection followed by clustering to estimate sequences containing a simultaneous events. However, scaling EBM fixes the number of simultaneous events in each stage a priori and does not allow mixtures of serial and simultaneous events in a single sequence. P-EBM therefore offers a more flexible approach as it can estimate arbitrary arrangements of simultaneous events in a purely data-driven way. Finally, the subtype and stage inference (SuStaIn) model (Young et. al. 2018) generalises EBM by introducing a linear z-score model and sequence subtypes. As with the EBM, SuStaIn assumes that biomarker abnormalities (in this case z-score events) occur in serial. The SuStaIn algorithm can implement other EBM’s, such as P- EBM, as the base model, a promising avenue of research.

### 5.5. Future opportunities and challenges

The P-EBM may be applied to identify key stages of disease in real-world datasets. One practical application of P-EBM is to inform biomarker collection strategies:- the shorter sequences estimated by P-EBM suggest which biomarkers are needed for patient staging. Reducing biomarker collection requirements for patient staging has tangible economical and patient benefits. We show patient stages estimated from a reduced biomarker subset are highly consistency with those estimated on the full dataset. Furthermore, P-EBM may inform studies of disease mechanisms. The temporal sequence of biomarker events inferred by the P-EBM may suggest new hypotheses regarding the causes and effects of disease, which may be corroborated using mechanistic models of disease evolution or laboratory experiments.

Several challenges may be addressed in future work. Firstly, the P-EBM search space grows super-factorially with the number of biomarkers. Efficient optimisation methods will be beneficial when applying the P-EBM to very large datasets. Model complexity may be accounted for when comparing the P-EBM and EBM model fits, such as through validation on held-out data. Finally, the P-EBM assumes subjects are sampled from a single disease trajectory. Integrating P-EBM and SuStaIn to account for both multiple disease subtypes and simultaneous events may enable more parsimonious representations of disease heterogeneity (Young et. al. 2018).

## 6. Conclusion

The P-EBM generalises event-based modelling of disease progression to enable estimation of parsimonious sequences of disease progression where multi-biomarker events can occur simultaneously within a common stage. High fidelity was observed in a range of realistic simulation scenarios and P-EBM suggested reduced biomarker subsets for patient staging. In sporadic AD, P-EBM inferred a relatively short sequence which captures the main stages of AD progression and is consistent with known pathological changes. The P-EBM has potential to inform understanding of disease evolution and suggest more efficient biomarker collection strategies for patient staging in clinical trials and beyond.

## 7. Data and code availability

ADNI data is available online (adni.loni.usc.edu). P-EBM is available at https://github.com/csparker/pebm. Simulation data is available on request.

## 8. Author contributions

*CSP:* Conceptualisation, Methodology, Software, Formal analysis, Investigation, Writing - original draft, Writing – review & editing, Visualisation, Funding acquisition. *NPO:* Software, Resources, Data curation, Writing – review & editing, Supervision, Project administration, Funding acquisition. *ALY:* Software, Resources, Data curation, Writing – review & editing. *DCA:* Conceptualisation, Resources, Writing – review & editing, Funding acquisition. *GZ:* Conceptualisation, Methodology, Resources, Writing – review & editing, Supervision, Project administration, Funding acquisition.

