## Supplementary Tables and Figures for "Parsimonious EBM: generalising the event-based model of disease progression for simultaneous events"

### Supplementary Information

*S.1. Sequence perturbations for the simultaneous event-based model*

| Biomarker | Perturbed sequence |
|---|---|
| 1 | ({1}, {2}, {3}, {4}) |
| | ({1,2}, {3}, {4}) |
| | ({2}, {1}, {3}, {4}) |
| | ({2}, {3}, {1}, {4}) |
| | ({2}, {3}, {1, 4}) |
| | ({2}, {3}, {4}, {1}) |
| 2 | ({1, 2, 3}, {4}) |
| | ({1, 3}, {2}, {4}) |
| | ({1, 3}, {2, 4}) |
| | ({1, 3}, {4}, {2}) |
| 3 | ({3}, {2}, {1}, {4}) |
| | ({2, 3}, {1}, {4}) |
| | ({2}, {3}, {1}, {4}) |
| | ({2}, {1}, {3}, {4}) |
| | ({2}, {1}, {3, 4}) |
| | ({2}, {1}, {4}, {3}) |
| 4 | ({4}, {2}, {1, 3}) |
| | ({2, 4}, {1, 3}) |
| | ({2}, {4}, {1, 3}) |
| | ({2}, {1, 3, 4}) |

**Table S1.** Perturbations of the P-EBM sequence ({2}, {1, 3}, {4}). To generate each perturbation, a biomarker (left column) is chosen at random and replaced at a random valid position that results in a new sequence (right column).

*S.2. Subject demographics*

| Demographics | CN | MCI | AD |
|---|---|---|---|
| Number of subjects | 100 | 150 | 75 |
| Sex (M/F, %F) | 51/49 (51.0%) | 98/52 (65.3%) | 41/34 (54.6%) |
| Age (years, mean ± S.D) | 75 ± 5 | 73 ± 7 | 75 ± 8 |
| Edu (years, mean ± S.D ) | 15.7 ± 2.9 | 15.7 ± 3 | 15.1 ± 3 |
| APOE4 (+/-, %+) | 22/78 (22.0%) | 83/65 (56.1%) | 52/23 (69.3%) |

**Table S2.** Demographics of study participants from the ADNI dataset. Abbreviations: AD: Alzheimer's disease; APOE4: Apolipoprotein E gene; CN: Cognitively Normal; Edu: Education; M: Male; MCI: Mild Cognitive Impairment; F: female; S.D. standard deviation.

| Biomarker | Effect size | Simulation s.d. |
|---|---|---|
| Total Tau | 1.8 | 0.55 |
| Phosphorylated Tau | 2.4 | 0.42 |
| Amyloid-$\beta_{1-42}$ | 3.8 | 0.26 |
| Hippocampal volume | 2.0 | 0.50 |
| Entorhinal volume | 2.0 | 0.50 |
| ADAS-Cog | 3.2 | 0.31 |
| RAVLT | 2.7 | 0.37 |
| MMSE | 3.8 | 0.26 |
| Mid-temporal volume | 1.5 | 0.66 |
| Fusiform volume | 1.3 | 0.77 |
| Brain volume | 1.2 | 0.83 |
| Ventricle volume | 1.7 | 0.59 |

**Table S3.** Effect sizes and corresponding simulation standard deviations for biomarkers of the AD dataset.

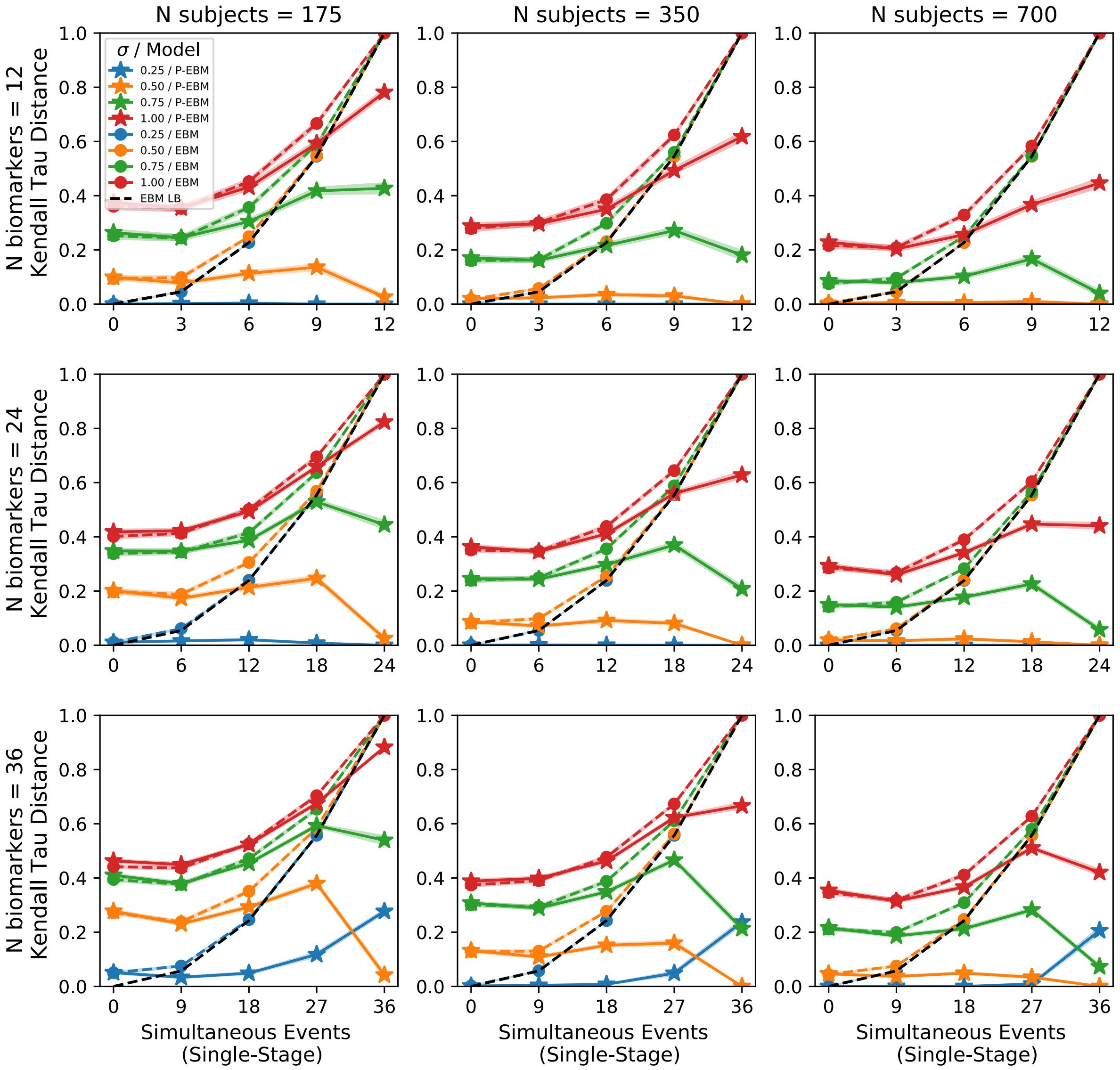


**Figure S1.** Kendall tau distance of the estimated sequences for ground truth sequences generated with a single stage containing a varying number of simultaneous events. Datasets were produced with a varying number of biomarkers (rows) and number of subjects (columns). Lines and shaded regions show the mean and standard error across the 100 simulated datasets. The dotted line shows the lower bound on the EBM distance metric.

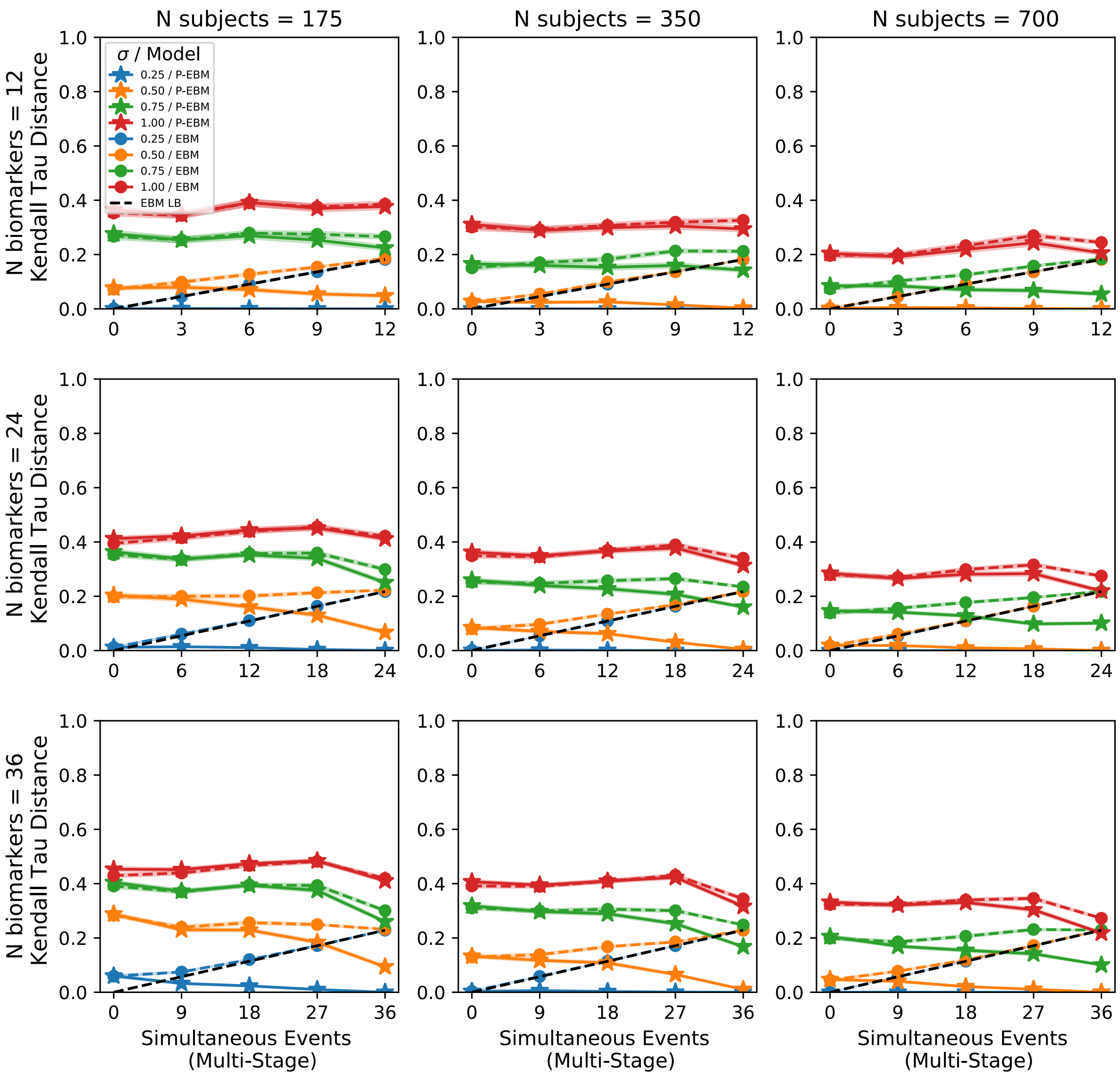


**Figure S2.** Kendall tau distance of the estimated sequences for ground truth sequences generated with multiple stages each with multiple simultaneous events. Datasets were produced with a varying number of biomarkers (rows) and number of subjects (columns). The number of simultaneous events in each stage is 3, 6 and 9 for datasets with 12, 24 and 36 biomarkers. Lines and shaded regions show the mean and standard error across the 100 simulated datasets. The dotted line shows the lower bound on the EBM distance metric.

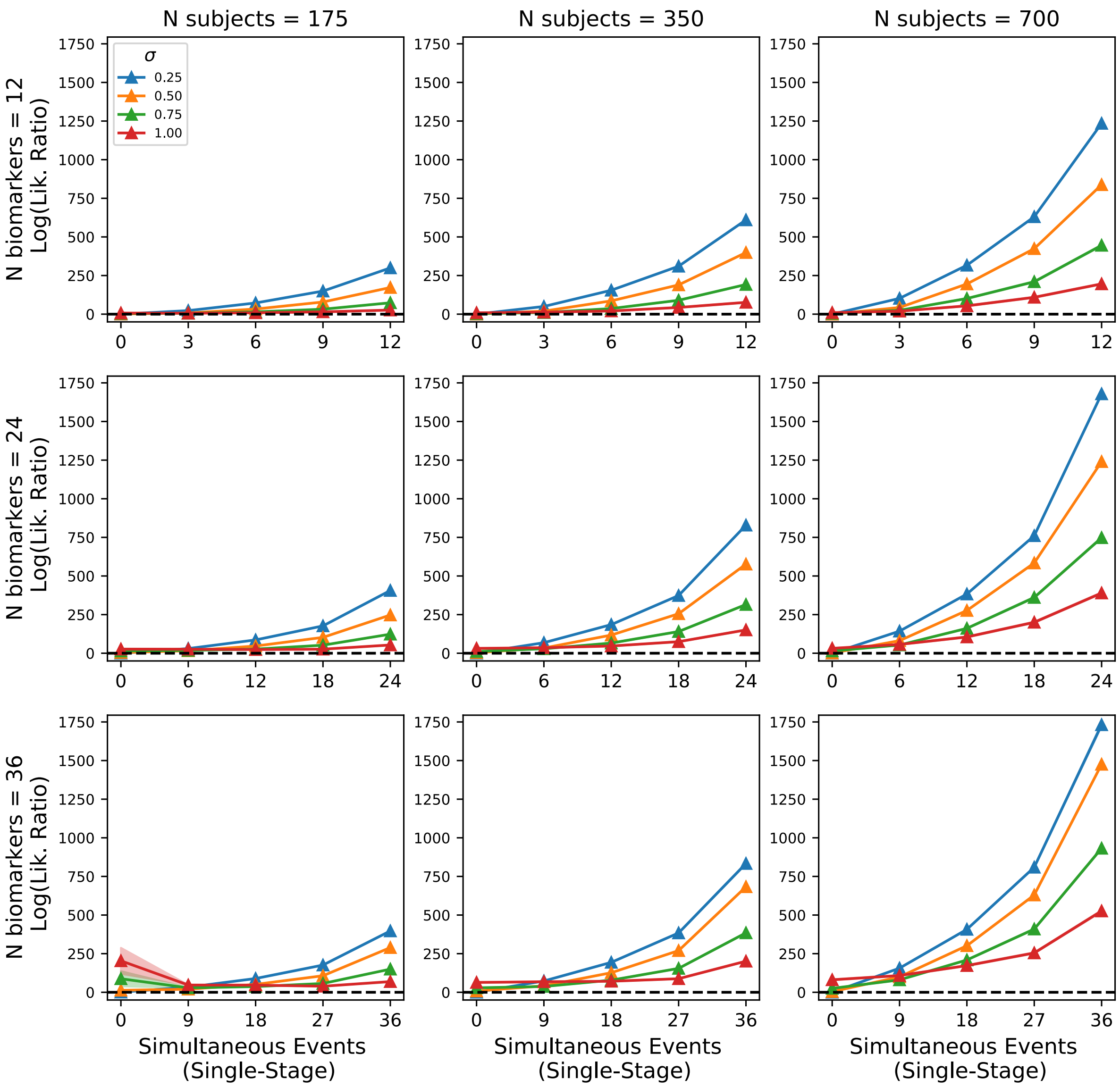


**Figure S3.** Log of the likelihood ratio between the P-EBM and EBM estimated sequence for ground truth sequences generated with a single stage containing a varying number of simultaneous events. Datasets were produced with a varying number of biomarkers (rows) and number of subjects (columns). Lines and shaded regions show the mean and standard error across the 100 simulated datasets.

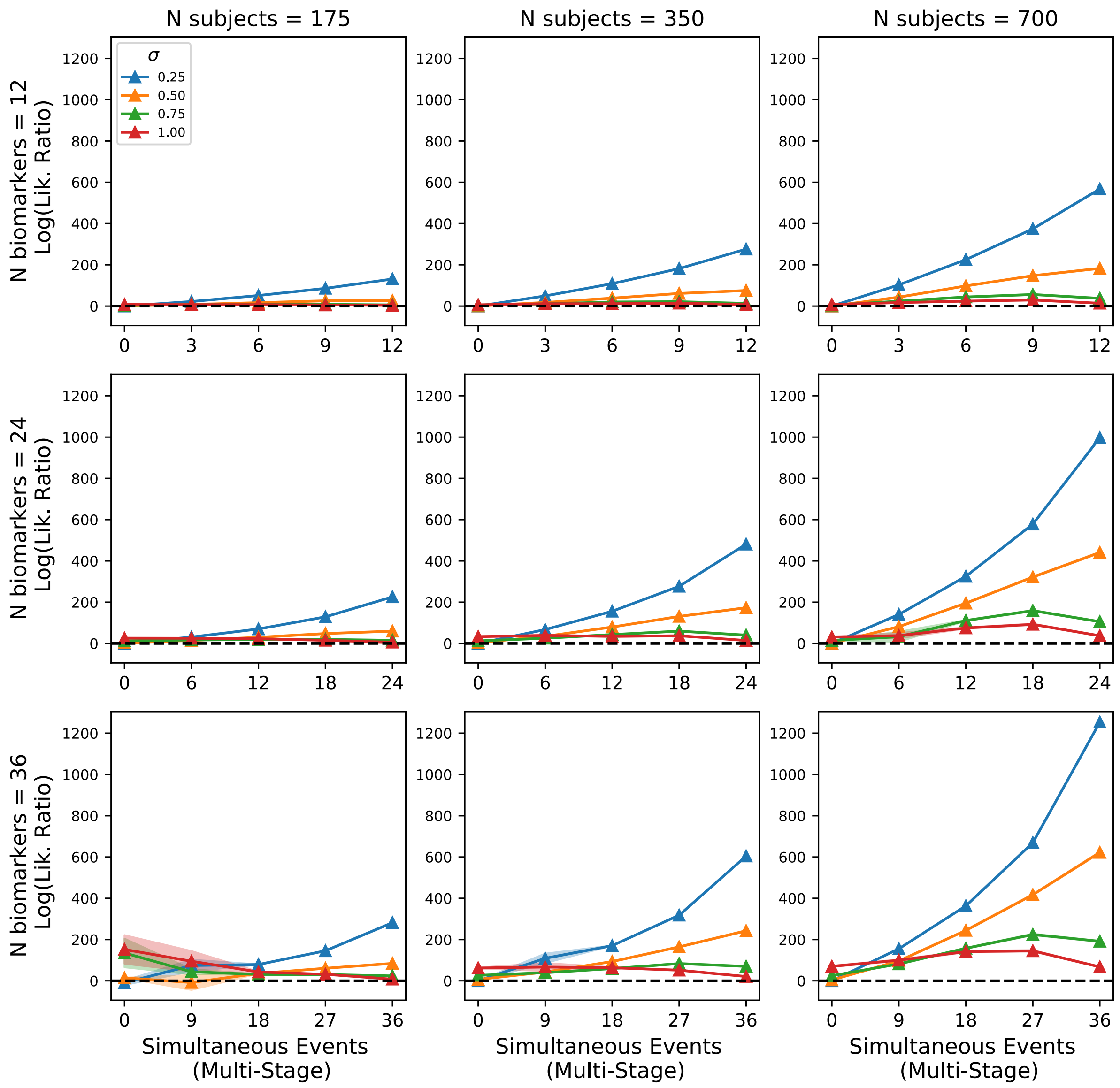


**Figure S4.** Log of the likelihood ratio between the P-EBM and EBM estimated sequence for ground truth sequences generated with multiple stages each with multiple simultaneous events. Datasets were produced with a varying number of biomarkers (rows) and number of subjects (columns). The number of simultaneous events in each stage is 3, 6 and 9 for datasets with 12, 24 and 36 biomarkers. Lines and shaded regions show the mean and standard error across the 100 simulated datasets.

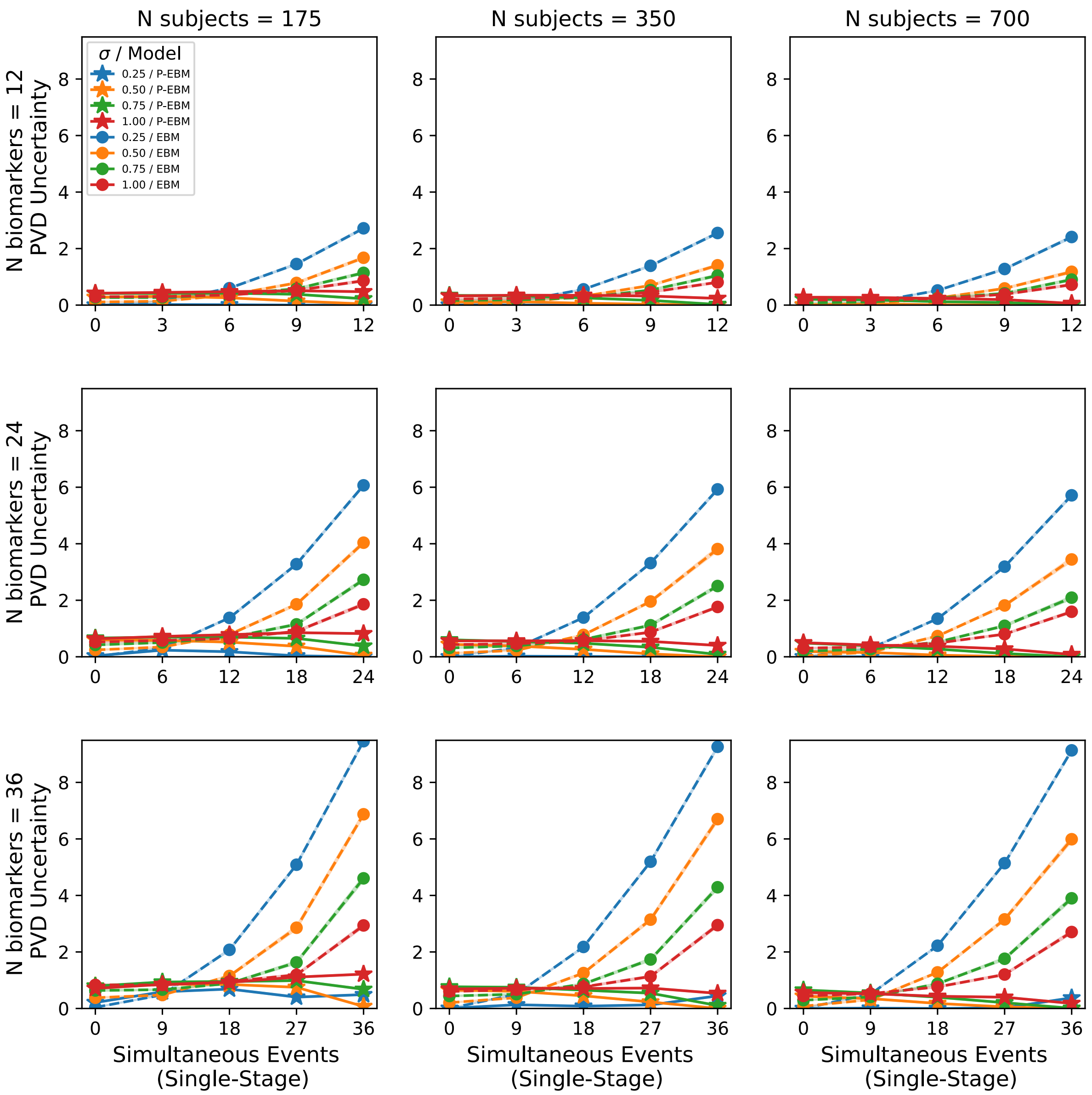


**Figure S5.** PVD uncertainty of the estimated sequence for ground truth sequences generated with a single stage containing a varying number of simultaneous events. Datasets were produced with a varying number of biomarkers (rows) and number of subjects (columns). Lines and shaded regions show the mean and standard error across the 100 simulated datasets.

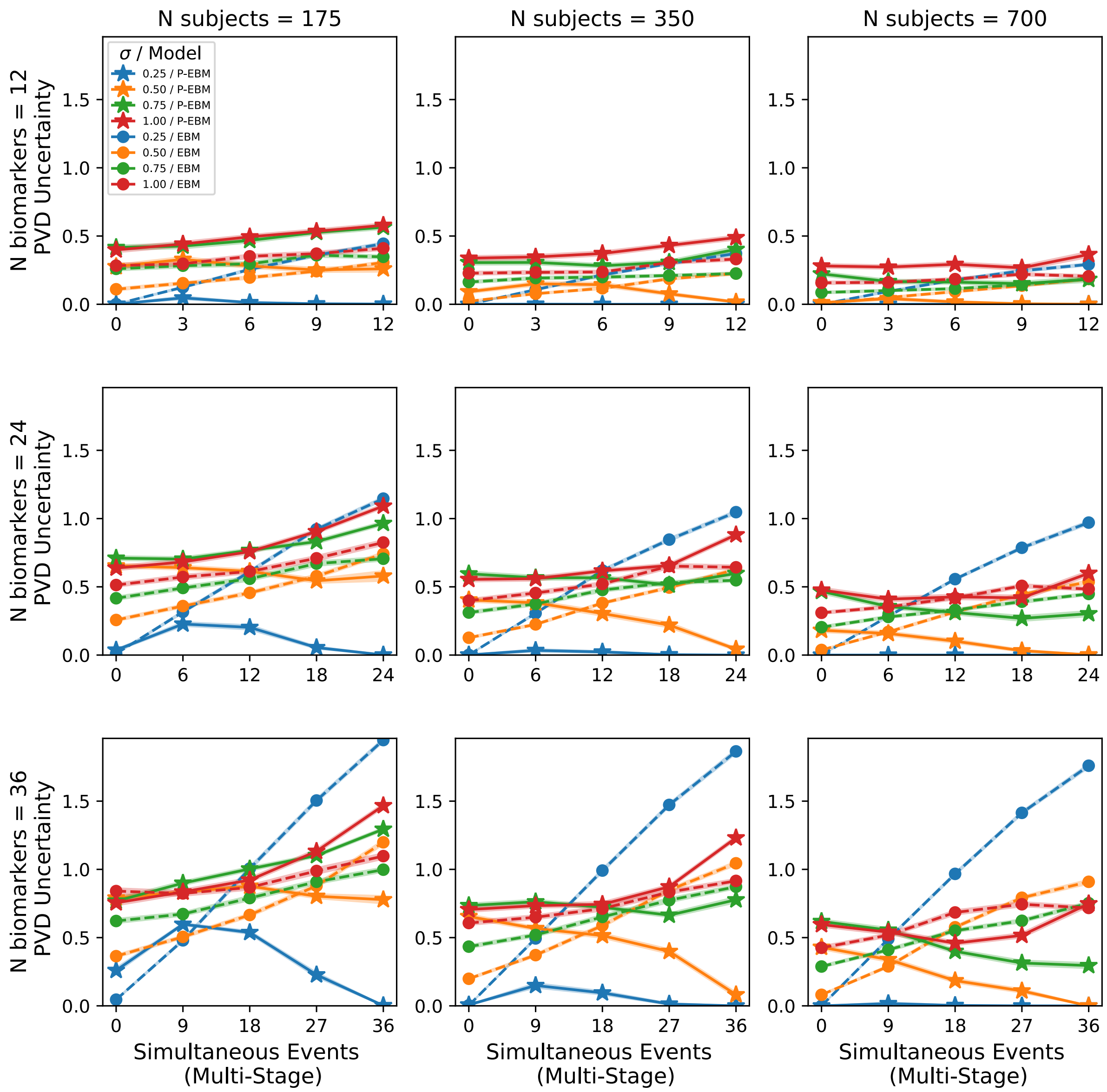


**Figure S6.** PVD uncertainty of the estimated sequence for ground truth sequences generated with multiple stages each with multiple simultaneous events. Datasets were produced with a varying number of biomarkers (rows) and number of subjects (columns). The number of simultaneous events in each stage is 3, 6 and 9 for datasets with 12, 24 and 36 biomarkers. Lines and shaded regions show the mean and standard error across the 100 simulated datasets.

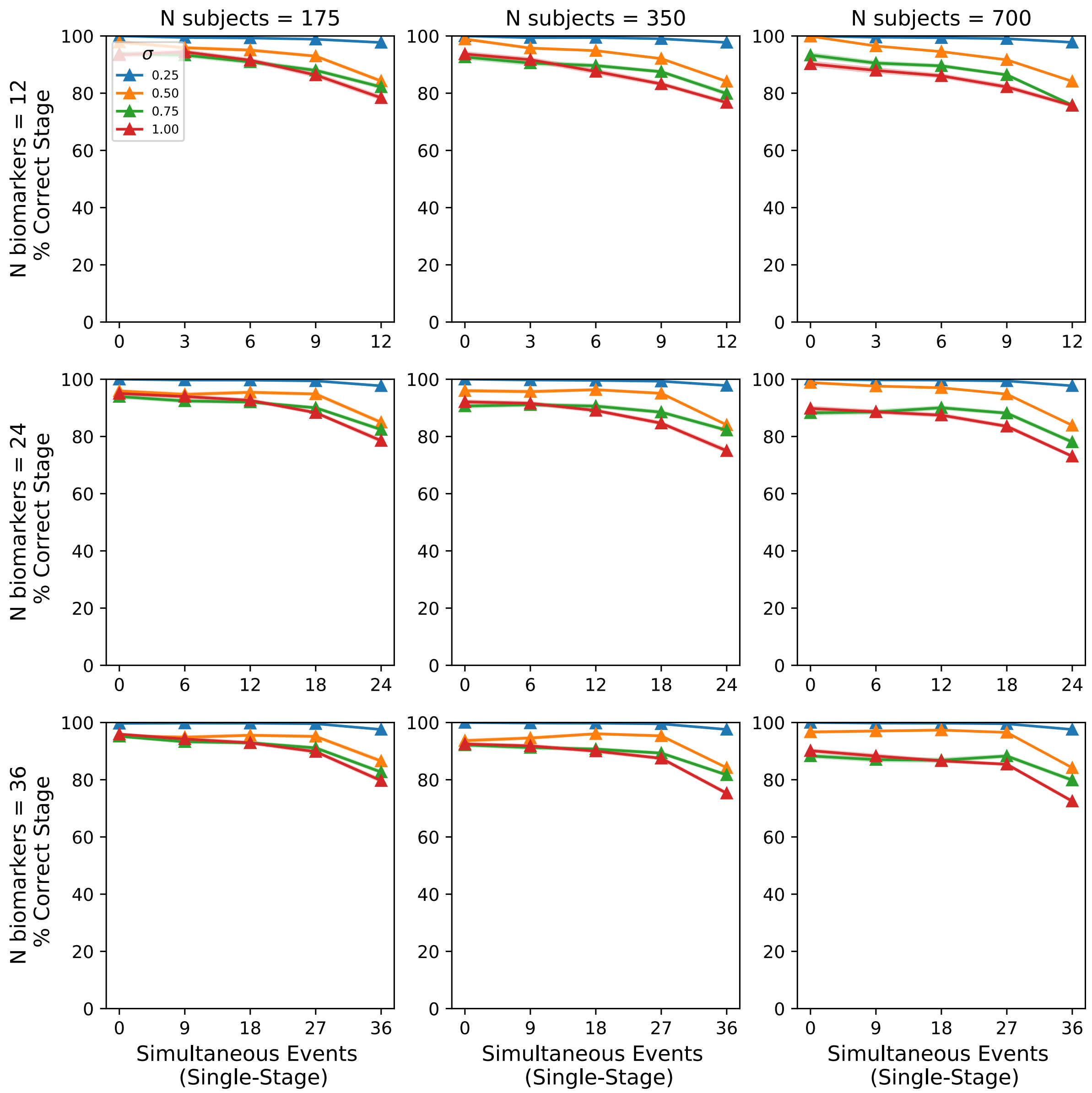


**Figure S7.** Percentage of subjects whose stage matches the full dataset when using the reduced dataset, for sequences generated with a single stage containing a varying number of simultaneous events. Datasets were produced with a varying number of biomarkers (rows) and number of subjects (columns). Lines and shaded regions show the mean and standard error across the 100 simulated datasets.

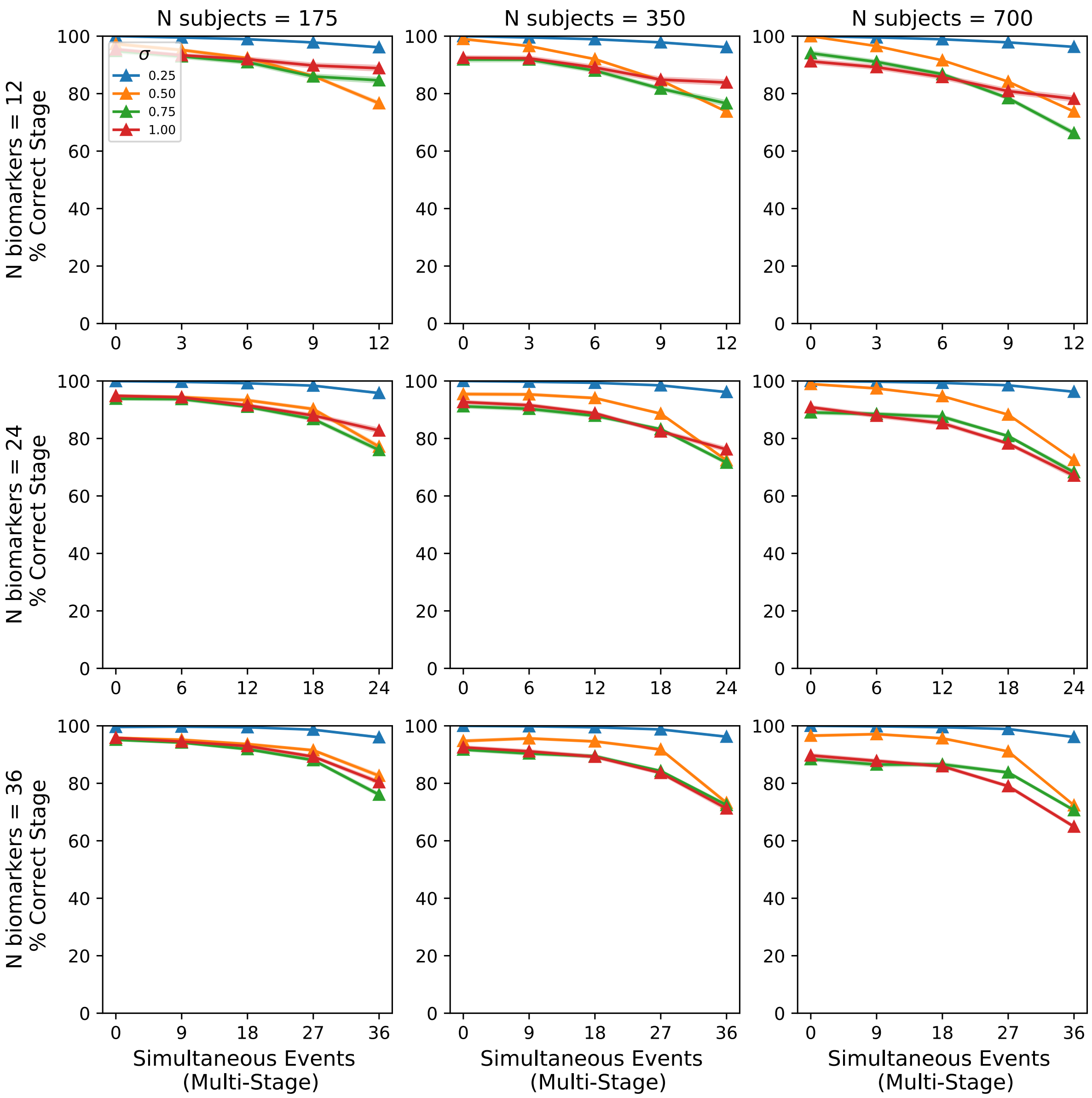


**Figure S8.** Percentage of subjects whose stage matches the full dataset when using the reduced dataset, for ground truth sequences generated with multiple stages each with multiple simultaneous events. Datasets were produced with a varying number of biomarkers (rows) and number of subjects (columns). The number of simultaneous events in each stage is 3, 6 and 9 for datasets with 12, 24 and 36 biomarkers. Lines and shaded regions show the mean and standard error across the 100 simulated datasets.

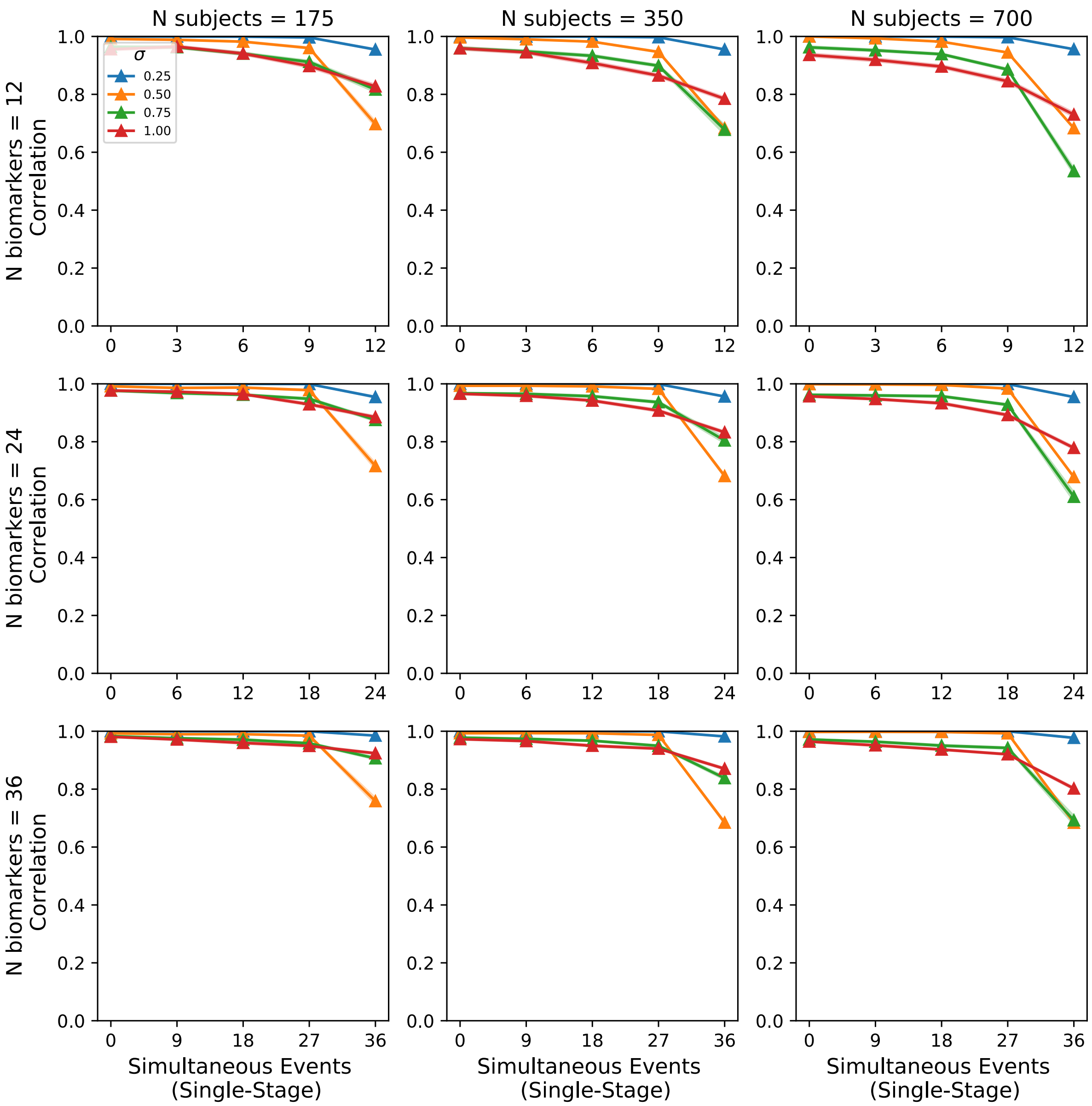


**Figure S9.** Correlation between the stage estimated on the reduced data and full data for sequences generated with a single stage containing a varying number of simultaneous events. Datasets were produced with a varying number of biomarkers (rows) and number of subjects (columns). Lines and shaded regions show the mean and standard error across the 100 simulated datasets.

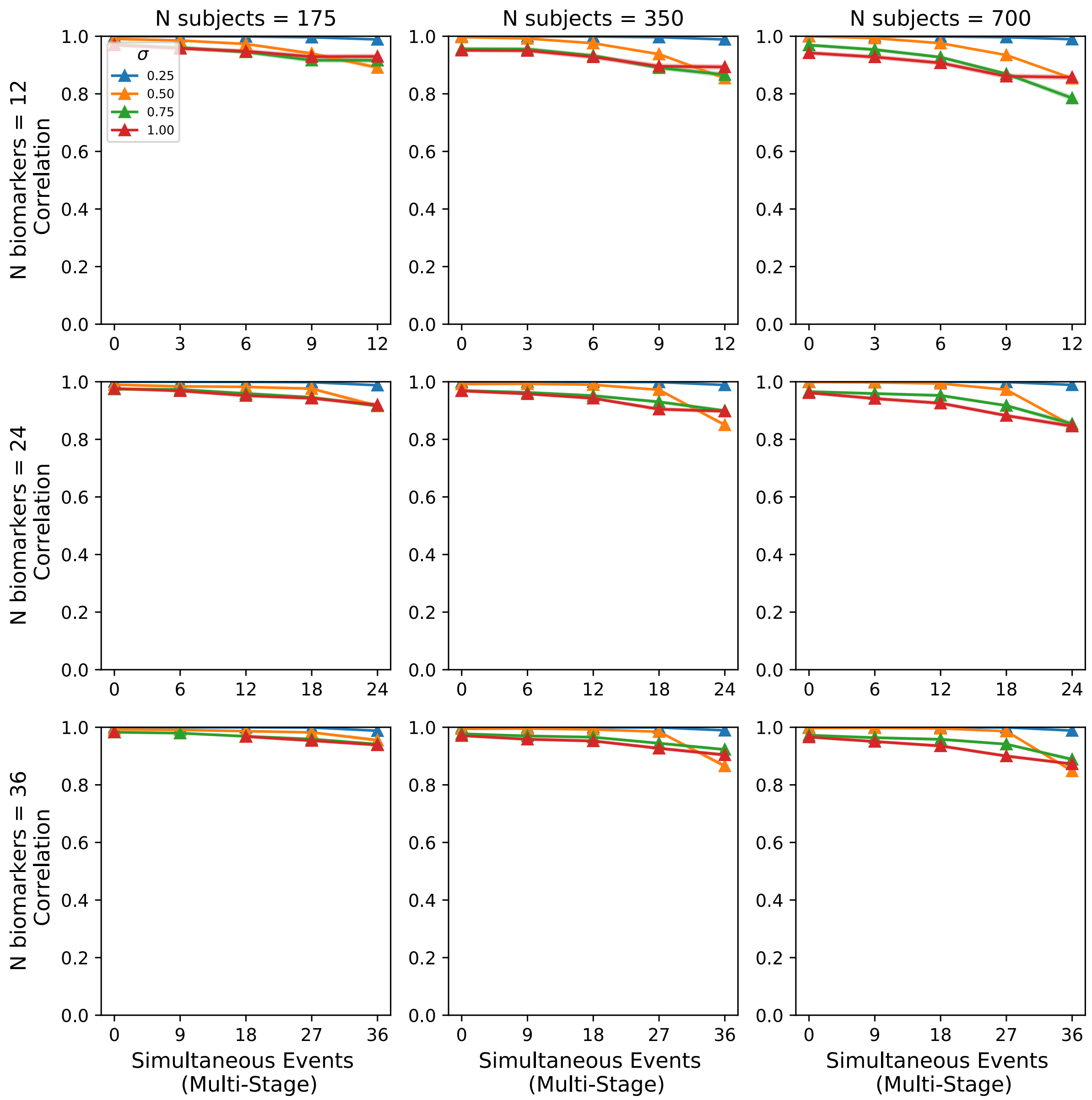


**Figure S10.** Correlation between the stage estimated on the reduced data and full data for sequences generated with multiple stages each with multiple simultaneous events. Datasets were produced with a varying number of biomarkers (rows) and number of subjects (columns). Lines and shaded regions show the mean and standard error across the 100 simulated datasets.
